# Secretion of an Argonaute protein by a parasitic nematode and the evolution of its siRNA guides

**DOI:** 10.1101/343772

**Authors:** FWN Chow, G Koutsovoulos, C Ovando-Vázquez, K Neophytou, JR Bermúdez-Barrientos, DR Laetsch, E Robertson, S Kumar, JM Claycomb, M Blaxter, C Abreu-Goodger, AH Buck

**Author notes:** Author’s contributed equally (FWN Chow, G Koutsovoulos, C Ovando-Vázquez). Current address: INRA, UMR 1355 Institute Sophia Agrobiotech, 06903 Sophia Antipolis, France. Current address: CONACYT-CNS-IPICYT, San Luis Potosí, SLP 78216, México.

## Abstract

Extracellular RNA has been proposed to mediate communication between cells and organisms however relatively little is understood regarding how specific sequences are selected for export. Here we describe a specific Argonaute protein (exWAGO) that is secreted in extracellular vesicles (EVs) released by the gastrointestinal nematode *Heligmosomoides bakeri*, at multiple copies per EV. Phylogenetic and gene expression analyses demonstrate exWAGO orthologues are highly conserved and abundantly expressed in related parasites but highly diverged in free-living genus *Caenorhabditis*. We show that the most abundant small RNAs released from the nematode parasite are not microRNAs as previously thought, but rather secondary small interfering RNAs (siRNAs) that are produced by RNA-dependent RNA Polymerases. The siRNAs that are released in EVs have distinct evolutionary properties compared to those resident in free-living or parasitic nematodes. Immunoprecipitation of exWAGO demonstrates that it specifically associates with siRNAs from transposons and newly evolved repetitive elements that are packaged in EVs and released into the host environment. Together this work demonstrates molecular and evolutionary selectivity in the small RNA sequences that are released in EVs into the host environment and identifies a novel Argonaute protein as the mediator of this.

## INTRODUCTION

Small RNA-mediated gene regulatory mechanisms are used by cellular organisms and viruses to enable various aspects of their development, defence strategies and physiology (1). One deeply conserved class of small RNAs is microRNAs (miRNAs), which associate with an Argonaute of the AGO clade to mediate the inhibition of translation of target messenger RNAs. MiRNAs were discovered for their crucial roles in development and are now appreciated to regulate numerous aspects of physiology and signalling. In the last 10 years, studies across a broad range of animal systems have implicated miRNAs in intercellular communication through their transport in extracellular vesicles (EVs) (2). Several reports also suggest mammalian miRNAs can move into other organisms, influencing gene expression and growth of microbes in the gut (3) and malaria parasites in the blood (4). We previously reported that the nematode parasite *Heligmosomoides bakeri* (renamed from *Heligmosomoides polygyrus* (5)) releases its own miRNAs within extracellular vesicles (EVs) and the EVs are internalized by mouse cells and suppress innate immune responses (6).

There is emerging data therefore in mammalian systems that suggest miRNAs can move between cells and organisms as a communication mechanism and that some pathogens have evolved to exploit this. However, despite these intriguing findings, there is very little understanding of how this is specified: do certain miRNAs get exported through active mechanisms or are the extracellular miRNAs a nonselected representation of the cytoplasm? Reports from different laboratories support both models, and indeed results may differ according to the cellular context, reviewed in (7). At the same time, mechanisms of specificity have begun to emerge from studies in several mammalian cell culture systems, where specific RNA-binding proteins recognize motifs in a subset of mature miRNAs and dictate their sorting into EVs (8-10).

We previously showed that the miRNAs released by the nematode parasite in EVs were a subset of those expressed in the adult worm (6). Here we sought to examine whether a parasite RNA binding protein could be associated with EV-RNA specificity. Our proteomic analysis identified one specific Argonaute protein in the excretory-secretory products and EVs of *H. bakeri* (6). Although there is controversy in whether Argonaute proteins are present in EVs in mammalian systems (7), this Argonaute was highly distinct from the ancestral Argonaute, belonging to a clade that has evolved specifically in nematodes (i.e. worm Argonautes or ‘WAGOs’) (11).

Rather little is known about WAGOs in nematode parasites but the free-living model organism *Caenorhabditis elegans* has at least four types of endogenous sRNAs and 25 Argonaute genes (12). In addition to miRNAs and piRNAs, *C. elegans* produces small interfering RNAs (siRNAs) from exogenous or endogenous double-stranded RNAs (dsRNAs). There is also a mechanism for *de novo* generation of siRNAs by RNA-dependent RNA polymerases (RdRPs), which are recruited to sRNA-target transcripts to amplify the silencing signal through the generation of secondary siRNAs. The secondary siRNAs dominate the sRNA content of adult *C. elegans* and have also been documented in several parasitic nematode species (13-16).

Here we examine the molecular and evolutionary properties of exWAGO in parasitic and free-living nematodes and demonstrate that exWAGO mediates the selective export of specific siRNAs in EVs. Unexpectedly, these siRNAs are more dominant in the EVs than miRNAs. We compare the genomic origin of siRNAs exported in EVs by *H. bakeri* to the resident siRNAs expressed in adults of both *H. bakeri* and *C. elegans*. Our results support a model where the resident sRNAs are dominated by secondary siRNAs, which are used for endogenous gene regulation and control of retrotransposons. In contrast, the parasite preferentially exports secondary siRNAs that are produced from newly evolved repetitive elements in the genome that associate with exWAGO. This adds evolutionary breadth to the handful of reports in mammalian systems suggesting RNA-binding proteins are a mechanism for selective RNA export and establishes *H. bakeri* as a tractable model for studying extracellular sRNA biology.

## METHODS

### Nematode lifecycle and sample collection

CBA x C57BL/6 F1 (CBF1) mice were infected with 400 L3 infective-stage *H. bakeri* larvae by gavage and adult nematodes were collected from the small intestine 14 days post infection. The nematodes were washed and maintained in serum-free media *in vitro* as described previously (17). For genomic analyses, DNA was collected from adult worms immediately following harvest from the gut, with extensive washing to remove host material followed by purification using Zymo Research Genomic DNA Clean & Concentrator kit following manufacturer’s instructions (further details in Supplementary Methods). RNA from *H.bakeri* was collected from adult worms in Qiazol (Qiagen) using mechanical disruption with 5 mm stainless steel beads (Qiagen) on a Tissue Lyser II (Qiagen). *C.elegans* were harvested and flash frozen as in (18). Total RNA was treated with RNA 5’ Polyphosphatase (Epicenter) following manufacturer’s instructions, before library preparation. Libraries for small RNA sequencing were prepared using the CleanTag small RNA library prep kit (Trilink) according to manufacturer’s instructions.

### Vesicle isolation and characterization

For collection of *H. bakeri* EVs described here, culture media from the adult worms was collected from 24-92 hours post harvest from the mouse (the first 24 hours of culture media was excluded due to potential host contaminants) [15]. Eggs were removed by spinning at 400 g and media was then filtered through 0.22mm filter (Millipore) followed by ultracentrifugation at 100,000 g for 2h in polyallomer tubes at 4 °C in a SW40 rotor (Beckman Coulter). Pelleted material was washed two times in filtered PBS at 100,000 g for 2h and re-suspended in PBS. The pelleted materials were mixed with 1.5 mL 2.5 M sucrose solution, and overlaid with a linear sucrose gradient (2.0 M − 0.4 M sucrose in PBS). Gradients were centrifuged 18-20h at 192,000g in a SW40 rotor (k-factor 144.5) (Beckman Coulter, Brea, CA). For RNA extraction and small RNA library preparation, the two fractions with densities of 1.16 − 1.18 g/cm^3^ (as calculated from measured reflective index by refractometry) were pooled, diluted 10 times in PBS and centrifuged again at 192,000g for 90 min in a SW40 rotor (k-factor 144.5) prior to resuspension in PBS.

Proteinase K sensitivity experiments were performed using 1.5ug (protein) of gradient-purified EVs treated with 5 ug /mL Proteinase K (Epicentre) in the absence or presence of 0.05% Triton X-100 at 37°C for 30 min. For western blot analysis, a polyclonal exWAGO antibody was used (generated and purified against peptides TKQTKDDFPEQERK, Eurogentec) overnight at 4°C, followed by incubation with a IRDye 680RD Goat anti-Rabbit IgG (LI-COR) secondary antibody for 1 hr at room temperature. Odyssey (LI-COR Biosciences) was used for visualization. Quantification was carried by western blot analysis in comparison to known concentrations of recombinant full length exWAGO containing an N-terminal 3XFlag-His tag using Image Studio Lite (LI-COR) software. For vesicle quantification, NTA was carried out using NanoSight LM14 instrument (Malvern Instruments, Malvern, UK).

### Genome assembly and protein-coding annotation

The short-read Illumina data were assembled and gapfilled using Platanus (19), and scaffolded using transcriptome evidence with SCUBAT2 (Koutsovoulos G. SCUBAT2, https://github.com/GDKO/SCUBAT2). Long-read PacBio data (a total of 10.2 Gb from reads with an N50 of 9,411 bases) were used to further scaffold and gapfill the assembly with PBJelly (20). We used the BRAKER (21) pipeline to predict protein-coding genes using the RNA-Seq reads as evidence. We combined the BRAKER general feature format (gff) file and the transcriptome assembly within MAKER2 (22) to predict untranslated regions of transcripts (UTRs) and remove low quality gene predictions. Further details are provided in Supplementary Methods and Supplementary Table 1.

### Identification of *H. bakeri* exWAGO orthologues in other Rhabditomorpha

Loci orthologous to the *H. bakeri* secreted WAGO (exWAGO) were identified using BLAST (23) searches of the genome-derived proteomes of *Haemonchus contortus, Necator americanus* and *Pristionchus pacificus*. The gene model of each identified orthologue was then evaluated with RNA-Seq data from the relevant species, and corrected if necessary. Using an alignment of these four proteins, a custom hidden Markov model (HMM) was constructed to identify homologues in other nematode genomes using HMMer (24) within GenePS (Koutsovoulos G https://github.com/jgraveme/GenePS). The discovered gene models were corrected based on alignment of the HMM profile, *de novo* Augustus (25) prediction, the previous predicted gene models and RNA-Seq data if available. Orthologues in *Caenorhabditis* species were validated through analysis of reciprocal best BLAST matches. Protein sequences of exWAGO orthologues were aligned with MAFFT (26) and the alignment was analysed with PHYML (using the LG+G model) (27,28). Bootstrap support was calculated from 100 bootstrap replicates.

### Ortholog clustering

Protein sequences of 21 nematode species were retrieved from the sources specified in Supplementary Table 2. Protein clustering was carried out using OrthoFinder v1.1.4 (29) under the MCL inflation value of 3.0. Orthogroups were analysed using KinFin v1.0.3 (30) by providing functional annotation and the phylogenetic tree of the taxa. Further details are provided in Supplementary Methods.

### Processing of sRNA-seq data

For this publication, we sequenced and analysed a total of 14 new sRNA-seq libraries from *H. bakeri*. These consisted of: adult worms (3 standard, 3 with 5’ RNA polyphosphatase treatment), purified extracellular vesicles (EVs) by sucrose gradients (2 standard, 2 with 5’ RNA polyphosphatase), and we also included the input pellet and supernatant before sucrose purification (2 each with 5’ RNA polyphosphatase treatment). In parallel we also prepared and sequenced 6 new *C. elegans* sRNA-seq libraries from adult worms (3 standard and 3 with 5’ RNA polyphosphatase treatment). Before proceeding we checked all libraries for their quality with FastQC (http://www.bioinformatics.babraham.ac.uk/projects/fastqc/). Reaper (31) was used to remove the Illumina small RNA Adapter sequences, and PullSeq 1.0.2 (https://github.com/bcthomas/pullseq) to keep reads that were at least 16 nucleotides long. For mapping to the genome, the alignment component of ShortStack 3.8.3 (32), with parameters: --nostitch, --mismatches 2, --mmap u, -- bowtie_m 500 and --ranmax 500 was used. The reference genome for *H. bakeri* genome was the one described in this paper, and for *C. elegans* the c_elegans.PRJNA13758.WBPS7.genomic.fa.gz file from https://parasite.wormbase.org.

### Annotation of ncRNA producing regions and differential expression

All the sRNA-seq results for each genome were used to predict novel ncRNA producing regions or clusters. ShortStack 3.8.3 was used with the following parameters: --pad 10, --mincov 10, --dicermin 18, --dicermax 32, and the resulting clusters were split at exon-intron boundaries. In order to obtain a level of expression for each sRNA-producing cluster, mapped reads were counted using the findOverlaps function from GenomicRanges R package with parameters minoverlap=16 and ignore.strand=TRUE. Only reads between 20-25nt were counted. For differential expression analysis, data from each genome were analyzed separately using the edgeR package (33). Only clusters with at least 0.5 counts-per-million in at least 2 libraries were considered. Different strategies were used to define the monoP and polyP-enriched clusters, and are fully described in Supplementary Methods.

### Information Content of sRNA-producing clusters

An Information Content (IC) value was calculated for each cluster, based on the coverage entropy compared to a uniform coverage distribution (maximum entropy). IC was calculated using the *entropy* R package (34), with the formula *log2(length(y)-entropy(y))*, where *y* is the cluster coverage. Further details are provided in Supplementary Methods.

### Argonaute Immunoprecipitations & qRT-PCR

Immunoprecipitations of adult *H. bakeri* worm lysates or EVs were carried out with rat polyclonal anti-exWAGO antibody (raised against full length protein) or rat IgG (control) using protein L magnetic beads (Fisher Scientific). Equivalent amounts of the input, flow-through and the immunoprecipitated product were kept for RNA extraction using the miRNeasy Serum/Plasma Kit (Qiagen). Synthetic spiked-in of 0. 1pM RT4 was added to the Qiazol before RNA extraction as internal control and 5 uL of total RNA was used as input for reverse transcription reactions using the miScript RT II System (Qiagen) followed by QuantiTec SYBR Green PCR kit (Qiagen). The RNAs were eluted with 14ul RNase-free water. The small RNAs that were associated with *H. bakeri* exWAGO were analysed by qRT-PCR with primers and conditions described in Supplementary Methods. The rabbit polyclonal anti-exWAGO antibody (as described above) was used for the western blot analysis of exWAGO.

## RESULTS

### A nematode-specific extracellular Argonaute is within extracellular vesicles released from *H.bakeri* at several copies per EV

We previously identified an Argonaute protein in the excretory-secretory and EV products of *H. bakeri* based on proteomic analyses (6). Several studies in mammalian systems have similarly reported Argonautes associated with EVs, in some cases under specific signalling conditions (35). However, Argonautes have also been reported to be contaminants that co-purify with EVs (8). In these samples we can detect exWAGO in both ultracentrifuge pellet and supernatant with a ~18 fold enrichment in EVs when loading equivalent protein volumes on a western blot. (Supplementary Figure 1). In order to rigorously determine whether the exWAGO in the ultracentrifuge pellet exists within EVs, we used ultracentrifugation followed by flotation on a sucrose gradient for purification, quantification by nanoparticle tracking analysis and visualisation by transmission electron microscopy. As shown in Figure 1, the *H. bakeri* EVs had a density of 1.16-1.18 g/cm3 and co-purified with exWAGO. We further subjected the sucrose-purified EV fractions to proteinase K treatment and confirmed that the exWAGO was protected from degradation but became susceptible when the EVs were lysed with detergent (Figure 1D). We analysed a defined number of sucrose-gradient purified EVs by western blot in comparison to recombinant exWAGO and found that exWAGO was present at 3.4 ± 1.1 copies per EV (Figure 1E and Supplementary Figure 1).

**Figure 1:**
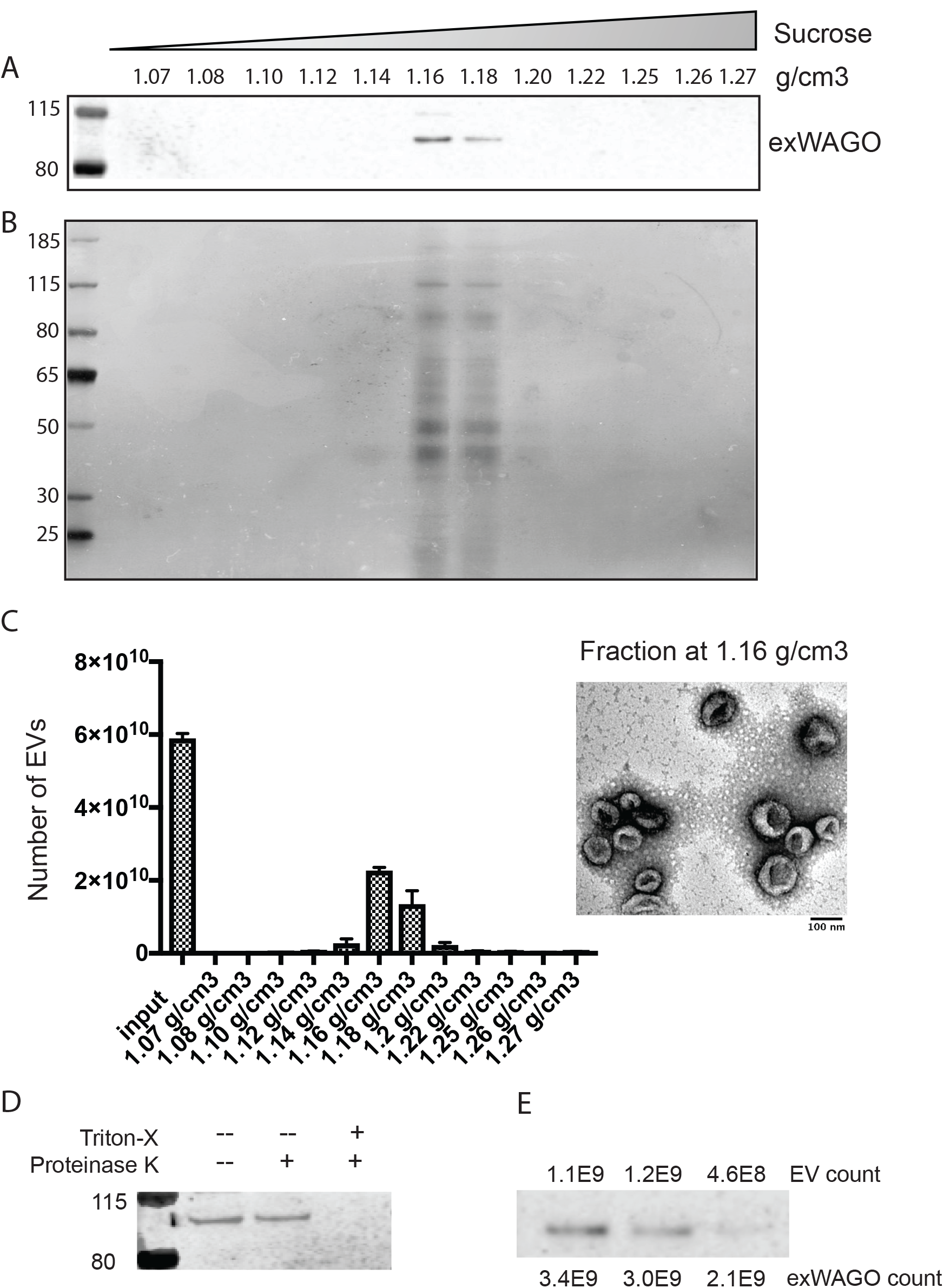
exWAGO is vesicular and present at multiple copies per EV. A) Western blot of equal volumes of sucrose-gradient fractions of EVs from *H. bakeri* using an antibody against exWAGO, B) Silver stain blot of same fractions, C) Nanoparticle tracking analysis of EV total number in each fraction (left) and TEM of 1.16 g/cm3 fraction (right). D) Western blot of exWAGO from gradient-purified EVs and following treatment with Proteinase K (5 ug/mL) with or without Triton-X (0.05%). E) Western blot of independent biological replicates of sucrose-gradient purified *H. bakeri* EVs, indicating number of EVs in each lane, quantified by NanoSight (Malvern). The exWAGO copy number was quantified in comparison to a standard curve based on recombinant exWAGO (shown in Supplementary Figure 1).

### An improved genome assembly and annotation to explore extracellular Argonautes and RNAs

In order to determine the full complement of Argonautes and small RNAs in *H. bakeri* we first generated a new genome assembly for this nematode based on combining short-read (~100-fold read coverage; Illumina) and long-read (~12-fold coverage; PacBio SMRT) data (Methods). The final genome assembly spans 697 Mb, 150 Mb longer than the first (Illumina-only) draft. While most sequenced nematode genomes are between 60 and 200 Mb, the strongylids (which include *H. bakeri*) tend to have larger genomes, ranging from 170 to 700 Mb (with a mean of ~380 Mb) (36). Our *H. bakeri* assembly is represented by 23,647 contigs (just over half the previous assembly’s 44,728 contigs), with an N50 of 180 kb (up from 36 kb). Assessment of genome completeness using the Core Eukaryotic Genes Mapping Approach (CEGMA) and Benchmarking Universal Single-Copy Orthologs (BUSCO) suggests ~88% of conserved genes are complete (~8% partial), with 96% of the assembled *H. bakeri* transcriptome mapping to the genome (Supplementary Table 1). Protein-coding genes were predicted with the BRAKER pipeline generating 23,471 protein-coding genes with 25,215 transcripts. Noncoding RNA genes, including rRNA, tRNA and miRNAs, were predicted using Rfam models and family-specific tools (see Supplementary Methods). The expansion of the *H. bakeri* genome compared to closely related clade V parasites is associated with an expanded repeat content. Over half (58.3%) of the *H. bakeri* genome contains some type of repeat element, including LINE elements (12.6% of the genome) and DNA elements (12.8%) (Table 1). Of all the repeats, 33.3% were found within genes (mostly in introns, which themselves occupy 33.5% of the genome). Interestingly, 30.6% of the genome was annotated as unclassified repeats, nearly two-thirds of which do not overlap any other kind of annotation.

**Table 1.** Non-overlapping genome composition.

|  | *C. elegans* |  | *H. bakeri* |  |
| --- | --- | --- | --- | --- |
|  | bases | % | bases | % |
| intergenic | 29,355,343 | 29.272 | 168,174,849 | 24.130 |
| exons | 28,409,938 | 28.329 | 23,682,169 | 3.398 |
| introns | 25,956,467 | 25.882 | 170,798,580 | 24.506 |
| transposons | 11,880,919 | 11.847 | 92,998,044 | 13.343 |
| novel repeats | 1,402,898 | 1.399 | 13,1452,375 | 18.861 |
| retroelements | 1,307,772 | 1.304 | 104,917,538 | 15.054 |
| satellite repeats | 594,478 | 0.593 | 377,947 | 0.054 |
| simple repeats | 581,880 | 0.580 | 2,960,599 | 0.425 |
| other ncRNA | 429,956 | 0.429 | 749,784 | 0.108 |
| piRNA | 256,574 | 0.256 | 63,990 | 0.009 |
| tRNA | 62,284 | 0.062 | 675,650 | 0.097 |
| miRNA | 35,312 | 0.035 | 43,955 | 0.006 |
| rRNA | 9,520 | 0.009 | 53,983 | 0.008 |
| yRNA | 3,060 | 0.003 | 5,940 | 0.001 |
| **TOTAL** | 100,286,401 | 100% | 696,955,403 | 100% |

### exWAGO is highly conserved and abundant in rhabditine (Clade V) parasitic nematodes and has diverged in *Caenorhabditis*

To determine the conservation of exWAGO across Clade V nematodes we clustered proteins from the new *H. bakeri* genome with proteomes predicted from the genomes of a selection of rhabditomorph nematodes (including *C. elegans* and six additional *Caenorhabditis* species, eleven strongyle parasites, the entomopathogen *H. bacteriophora*, the free living *Oscheius tipulae*, and the free-living diplogasteromorph *Pristionchus pacificus*) using OrthoFinder (29). The resulting orthogroups were interrogated with Kinfin (30) to identify orthologues of proteins predicted to be involved in RNAi in *H. bakeri* or known to be implicated in RNAi in *C. elegans*. This revealed that, as expected, nearly all of the machinery for miRNA and piRNA pathways, including highly conserved Argonautes ALG-1/2 and PRG1/2, is conserved across Clade V nematodes (Supplementary Figure 2). Strikingly, of the thirteen WAGOs in the *C. elegans* gene set, only four had coclustered orthologues from species other than *Caenorhabditis*. We therefore performed a joint phylogenetic analysis of all orthogroups containing Argonautes (defined by the presence of both PAZ and PIWI domains) (Figure 2A). This identified clades of Argonautes in parasitic species that were sister to *Caenorhabditis-specific* orthogroups. For example, the nuclear WAGOs HRDE-1 and NRDE-3 as well as WAGO-10 and WAGO-11 in *Caenorhabditis* are in fact orthologous to parasite-derived Argonautes in orthogroups OG01747 and OG07955 but their relationship has been obscured by differing rates of evolution in the different species groups. Similarly, the phylogenetic analysis shows that exWAGO does in fact co-cluster with a *Caenorhabditis-only* orthogroup that contains *C. elegans* SAGO-1, SAGO-2 and PPW-1. The orthogroup containing exWAGO contains Argonautes from many other parasitic strongyles, *H. bacteriophora, O. tipulae* and *P. pacificus* as well as Argonautes from *Caenorhabditis* species placed at the base of the genus (*C. monodelphis, C. castaneus*, and *C. sp*. 38) (Figure 2B). Examination of the intron-exon structure of these Argonautes supports this relationship (Figure 2C). The most basal *Caenorhabditis, C. monodelphis*, has a gene structure very similar to that of the other exWAGOs, but gene structure in other *Caenorhabditis* species has evolved rapidly. We suggest that *C. elegans* SAGO-1, SAGO-2 and PPW-1 are co-orthologues of *H. bakeri* exWAGO, and thus the biology of these genes may illuminate the origins and functions of exWAGO in parasites.

**Figure 2:**
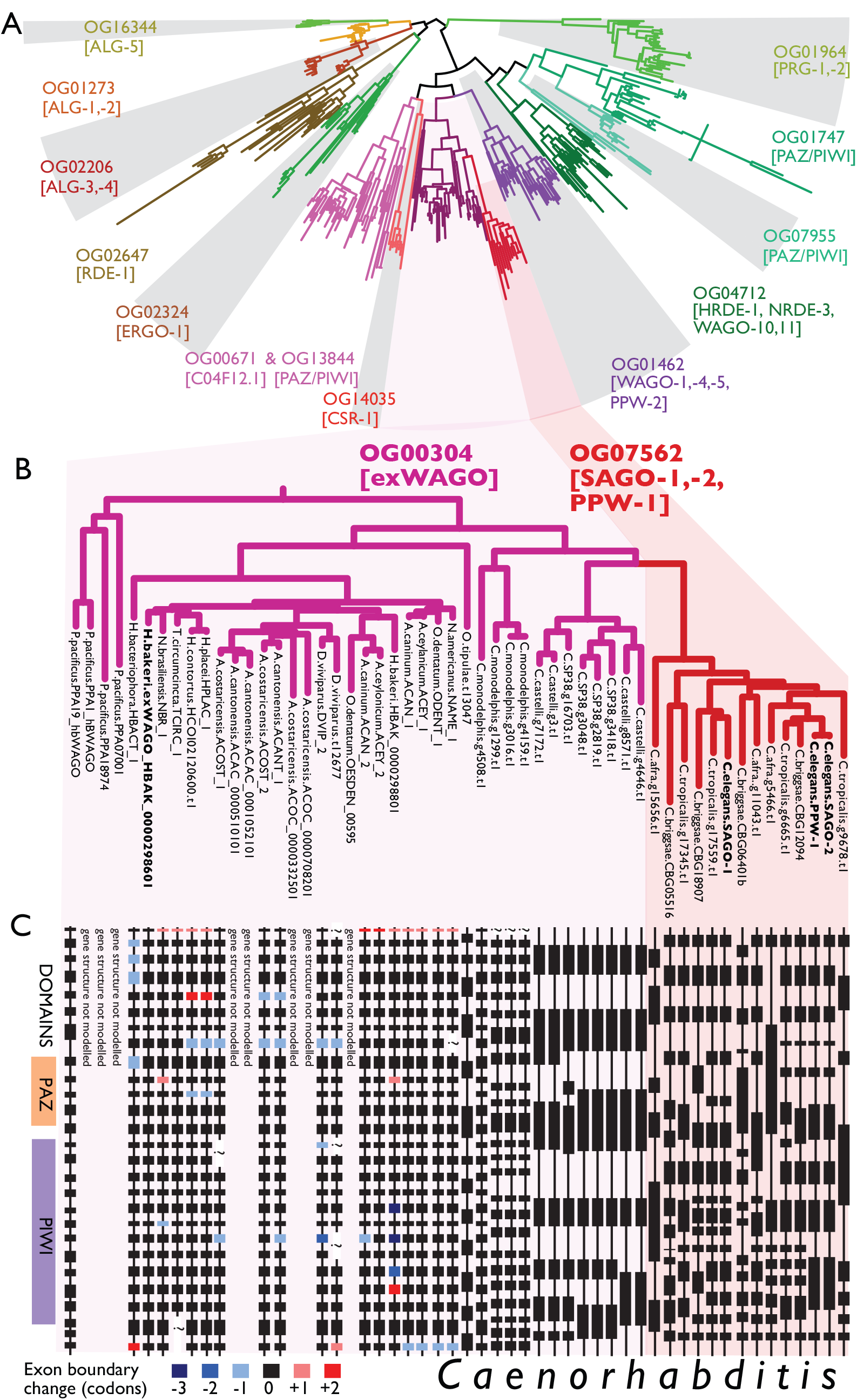
Phylogenetic tree of Argonautes in Clade V nematodes and gene structure of exWAGO. A) Grey shading denotes different orthogroups, *C. elegans* protein names in each clade are noted (or absent if no orthologues in that Clade). B) Tree showing phylogenetic relationship and branch lengths of exWAGO orthologues across Clade V, C) Conservation of exons and introns in exWAGO homologues. Each box is an exon with the width denoting length. Boxes with dashed lines denote exons with possible errors in the genome assembly of the species. Colours denote differences in exon size in triplets compared to exWAGO.

Using our new annotation of Argonautes we used existing RNAseq data to determine the expression levels of all Argonautes in adult life stages of parasitic versus free-living Clade V nematodes. Strikingly, we found that exWAGOs are the most abundantly expressed of all Argonautes (Supplementary Table 3) in the sheep parasites *Haemonchus contortus* and *Teladorsagia circumcincta* and the lungworm *Angiostrongylus cantonensis*. It is also the second most highly expressed in the human hookworm, *Necator americanus* (Figure 3). In contrast, the SAGO-1, 2 and PPW orthologs in *C. elegans* adults are not expressed at high levels (Supplementary Table 3). We further identified exWAGO in the excretory-secretory (ES) products of adult *Nippostrongylus brasiliensis* (another rodent parasite) (Table 2). No peptides mapping to any other Argonaute proteins were identified in multiple samples, pointing to a unique extracellular role for this particular Argonaute across the parasite species.

**Figure 3:**
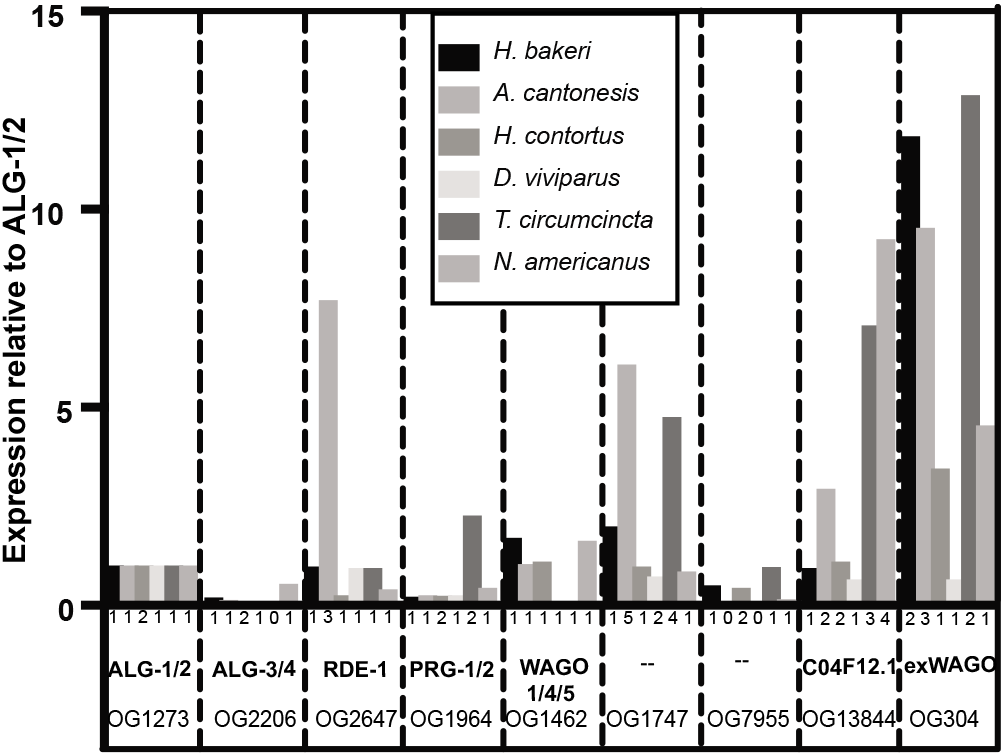
Expression of Argonautes across Clade V parasitic nematodes. Relative expression levels of Argonautes from RNAseq data of the adult parasites noted. Data were based on the sum of tpm reads for each orthogroup (defined in Figure 2), normalized to tpm for OG1273 orthogroup (ALG-1/2). The total number of distinct transcripts in each orthogroup in each species is noted below each column. The known *C.elegans* Argonaute names are used where applicable, or exWAGO as defined in this work.

**Table 2.**
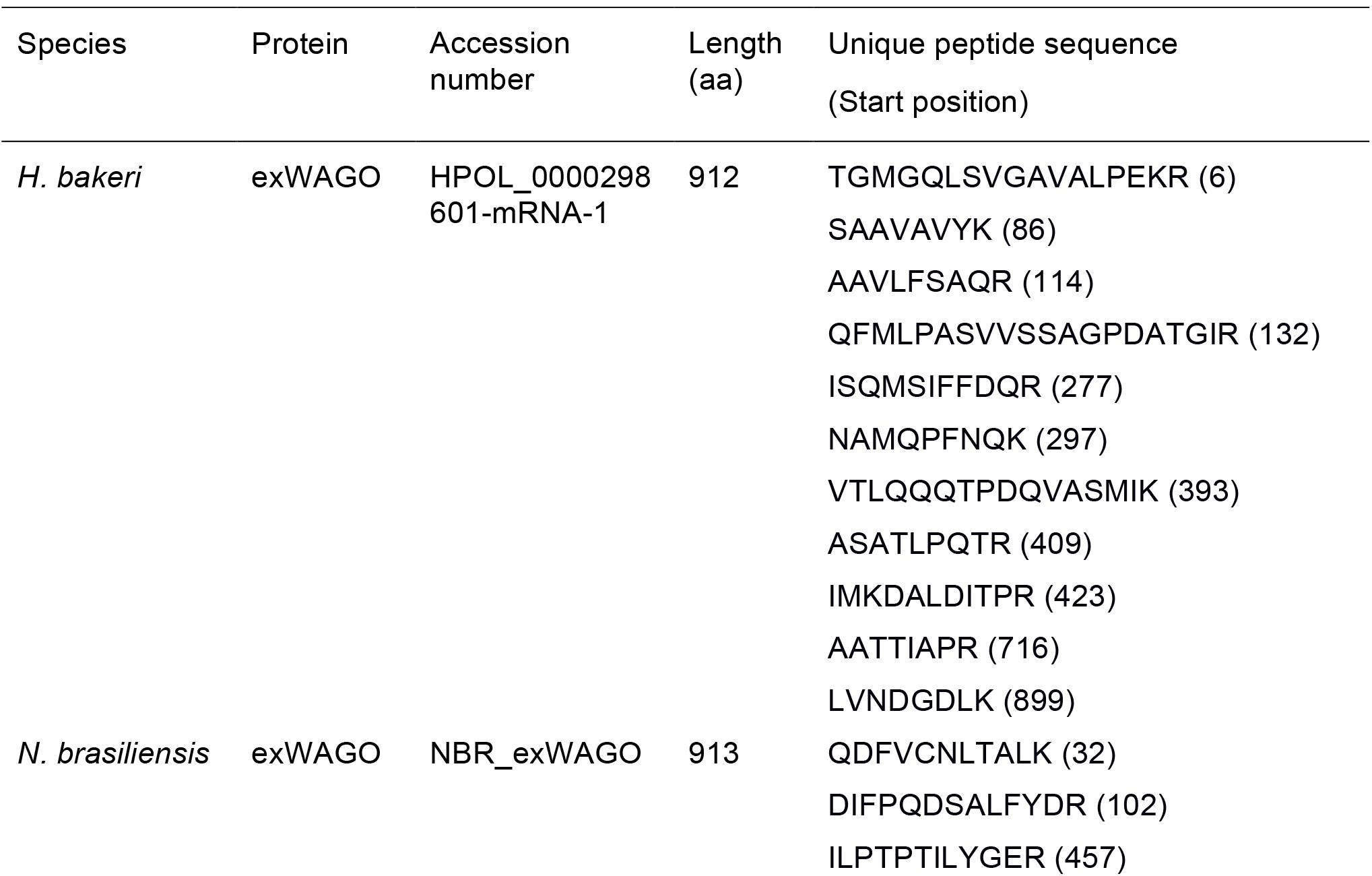
ExWAGO identified in EV products by mass spectrometry.

### Comparative analysis of resident sRNA distribution in *H. bakeri* versus *C. elegans* suggests shared functions in endogenous gene regulation

To understand whether the dominance of exWAGO in the parasites was reflected at the level of sRNA composition we first carried out side-by-side analysis of endogenous sRNAs present in *H. bakeri* and *C. elegans* adult nematodes. sRNA datasets were generated in triplicate, capturing either only 5’-monophosphate RNAs or all RNAs (after treatment with 5’ polyphosphatase). As expected, the untreated libraries from whole nematodes were dominated by reads mapping to miRNAs in each genome, having the characteristic first nucleotide preference of U and peak length of 22 nt (Figure 4A,D). In contrast, the whole-nematode libraries treated with 5’ polyphosphatase showed a clear enrichment for RNAs with a first base preference of guanine, the majority of which were 22 nt in length in *C. elegans* and 23 nt in *H. bakeri* (Figure 4B,E). This signature is characteristic of secondary siRNA products of RdRPs, and suggests secondary siRNAs dominate the resident sRNA populations of both nematodes. The length variation (22 *versus* 23 nt) may indicate mechanistic differences between the RdRPs that generate them or the Argonaute proteins that stabilize them.

**Figure 4:**
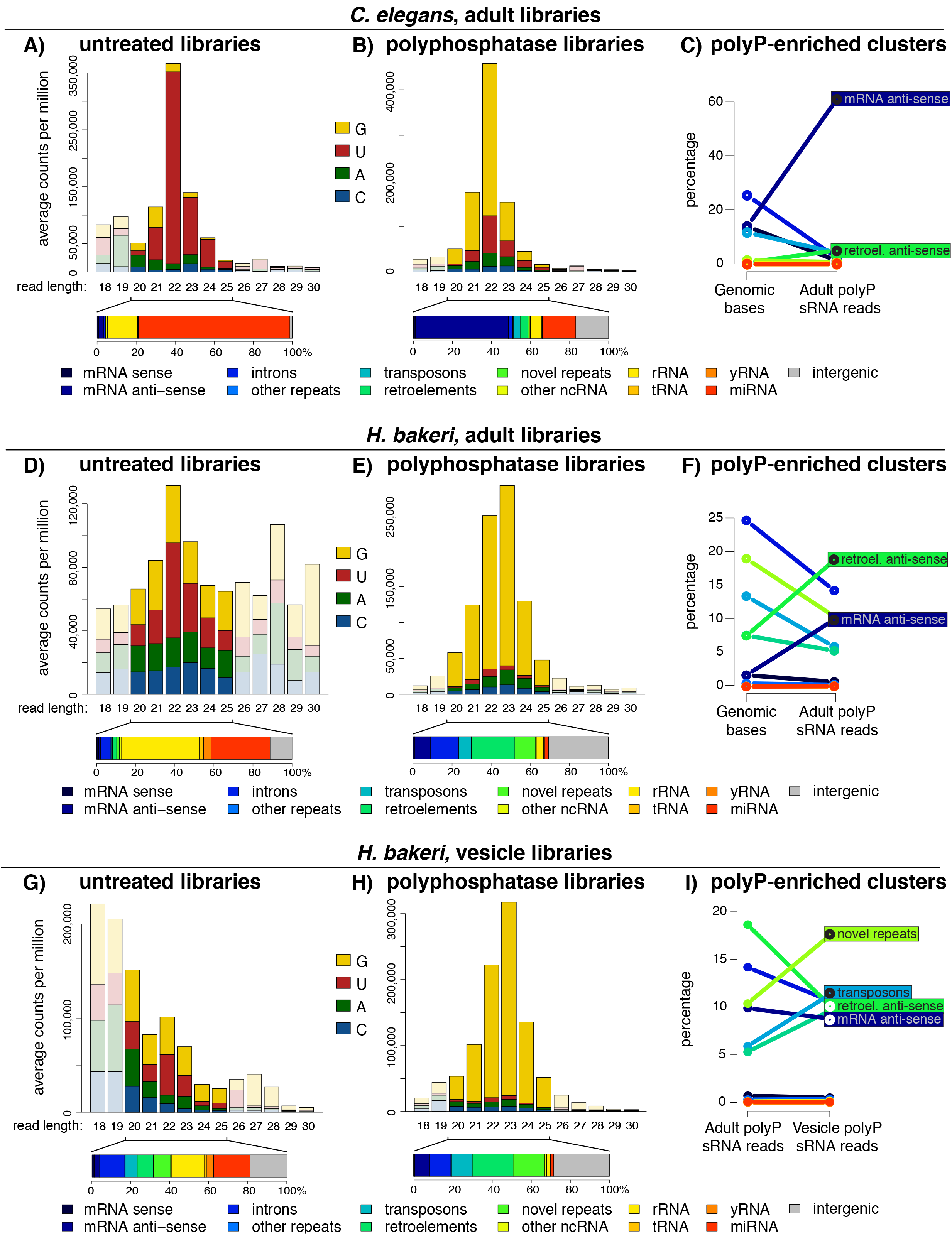
sRNA composition in adult *C. elegans* and *H. bakeri*, compared to *H. bakeri* extracellular vesicles. First nucleotide and length distribution of untreated small RNA libraries for *C. elegans* adults (A), *H. bakeri* adults (D) and *H. bakeri* EVs (G), and their corresponding polyphosphatase-treated libraries (B,E,H). The proportions of the 20-25 nt reads mapping within annotated categories in the genome (from Table 1) are shown beneath each barplot. Line plots for *C. elegans* (C) and *H. bakeri* (F) showing the relationship between the percentage of the genome occupied by each annotation category and the percentage of 20-25 nt reads from the polyP-enriched clusters, while (I) shows the relationship between the percentage of the 20-25 nt reads from the adult polyP-enriched clusters to those from the EV polyP-enriched clusters (see Supplementary Methods).

By comparing the 5’ polyphosphatase-treated and untreated libraries in both species, we identified 137,531 regions in the *H. bakeri* genome that have more mapped reads from the 5’ polyphosphatase-treated libraries (we call these regions polyP-enriched clusters) and 6,075 regions with relatively more reads from the untreated libraries (monoP-enriched clusters, see Supplementary Methods and Supplementary Figure 3). We reasoned that these represent sRNAs with two distinct modes of biogenesis, with the monoP-enriched clusters containing sRNAs cleaved by ribonucleases (such as Dicer) or being degradation products, and the polyP-enriched clusters containing unprocessed products of RdRPs or RNA polymerase III. Consistent with this model, the monoP-enriched clusters contained a higher fraction of miRNA-mapping reads than the untreated libraries, while the polyP-enriched clusters had a much reduced fraction of miRNA reads (Supplementary Figure 4). The same general strategy was applied to the *C. elegans* sRNAs, to compare the polyP-enriched clusters of both nematodes, which represent by far the most abundant type of sRNA in adults. The majority (62.3%) of the reads within the polyP-enriched clusters of *C. elegans*, mapped antisense to messenger RNAs, consistent with roles in regulating endogenous gene expression (Figure 4C). In contrast, 9.9% of the reads from polyP-enriched clusters of *H. bakeri* mapped antisense to mRNAs (Figure 4F). To compare these numbers, we need to take into account the fraction of each genome occupied by mRNAs. The *C. elegans* genome devotes 28.3% of its base pairs to coding exons (Table 1, Supplementary Figure 5), therefore if sRNAs were produced randomly across the genome 14.1% would map antisense to these. Consequently, our observed proportion of antisense mRNA polyP sRNAs represents a 4.4-fold increase over what is expected by chance. Since only 3.4% of the *H. bakeri* genome encodes exons, the polyP sRNAs that map antisense to mRNAs are 5.8-fold more frequent than expected. Following similar logic, both nematodes have a significant overrepresentation of polyP sRNAs mapping antisense to known retrotransposons (7.4-fold increase in *C. elegans*, 2.5-fold increase in *H. bakeri*, Figure 4C,F). Thus, despite the drastic differences in genome content of the two species, there is a conserved pattern of siRNAs that likely reflect common functionality in endogenous gene regulation and genome defence.

### Vesicular siRNAs are largely derived from novel-repeat elements and associate with exWAGO

Our previous work indicated that EVs secreted from *H. bakeri* adults are associated with a population of sRNAs. We characterized miRNAs and Y RNAs, but only sequenced sRNAs with a 5’ monophosphate (6). To generate a more comprehensive characterization of EV sRNAs and examine selectivity, we analysed duplicate sRNA datasets from purified EVs, capturing either only 5’- monophosphate RNAs or all RNAs (after treatment with 5’ polyphosphatase). To ensure the sequenced sRNAs were derived from EVs, and not co-purifying or free complexes, EVs were purified by ultracentrifugation and sucrose gradient prior to RNA extraction, library preparation and sequencing. We detected several species of miRNA in EV-derived libraries as expected from our previous work, however the vast majority of EV sRNAs are 23G siRNAs, only detected with 5’ polyphosphatase treatment (Figure 4H). To focus on the RdRP products, we selected the polyP-enriched clusters (Supplementary Figure 3). EV-derived, polyP-enriched sRNAs had a 1.9-fold enrichment for siRNAs derived from transposons and a 1.7-fold enrichment for species-specific repeats of the *H. bakeri* genome, compared to the polyP-enriched sRNAs from adults (Figure 4I). In contrast, the siRNAs mapping antisense to protein coding genes and retrotransposons were relatively depleted within the EV libraries. Because some exWAGO can be found outside of EVs, we also sequenced libraries from the supernatant of the ultracentrifuge pellets. Strikingly, the polyP-enriched sRNAs in the supernatant do not show the same pattern of enrichment for species-specific elements and cluster more closely with the libraries of the adults (Supplementary Figure 6). We also confirm by qRT-PCR that the levels of EV-enriched siRNAs are higher in the EVs than the supernatant (Supplementary Figure 6). These results suggest selective partitioning of siRNAs into the EVs and identifies recently evolved regions of the genome as a primary source of EV siRNA.

To further explore and quantify this selectivity, we calculated, for each polyP-enriched cluster, a measure of entropy-based Information Content (IC) using either adult or EV reads (see Methods). The higher the IC value, the more concentrated the reads are in a few peaks, while the lower the IC value, the more evenly distributed the reads are across the cluster (e.g. Figure 5A, inset). Interestingly, the IC values are consistently higher for reads coming from EVs than from adult libraries (Figure 5A), indicating that the EVs more often contain reads from specific peaks and are not a random sampling of the adult sRNA pool. Figure 5B illustrates a region in the *H. bakeri* genome that produces siRNAs enriched in the EVs. The only annotated elements in this region are repeats, mostly novel (species-specific) elements. These results support the idea that EV content reflects a selection of specific siRNA sequences.

**Figure 5:**
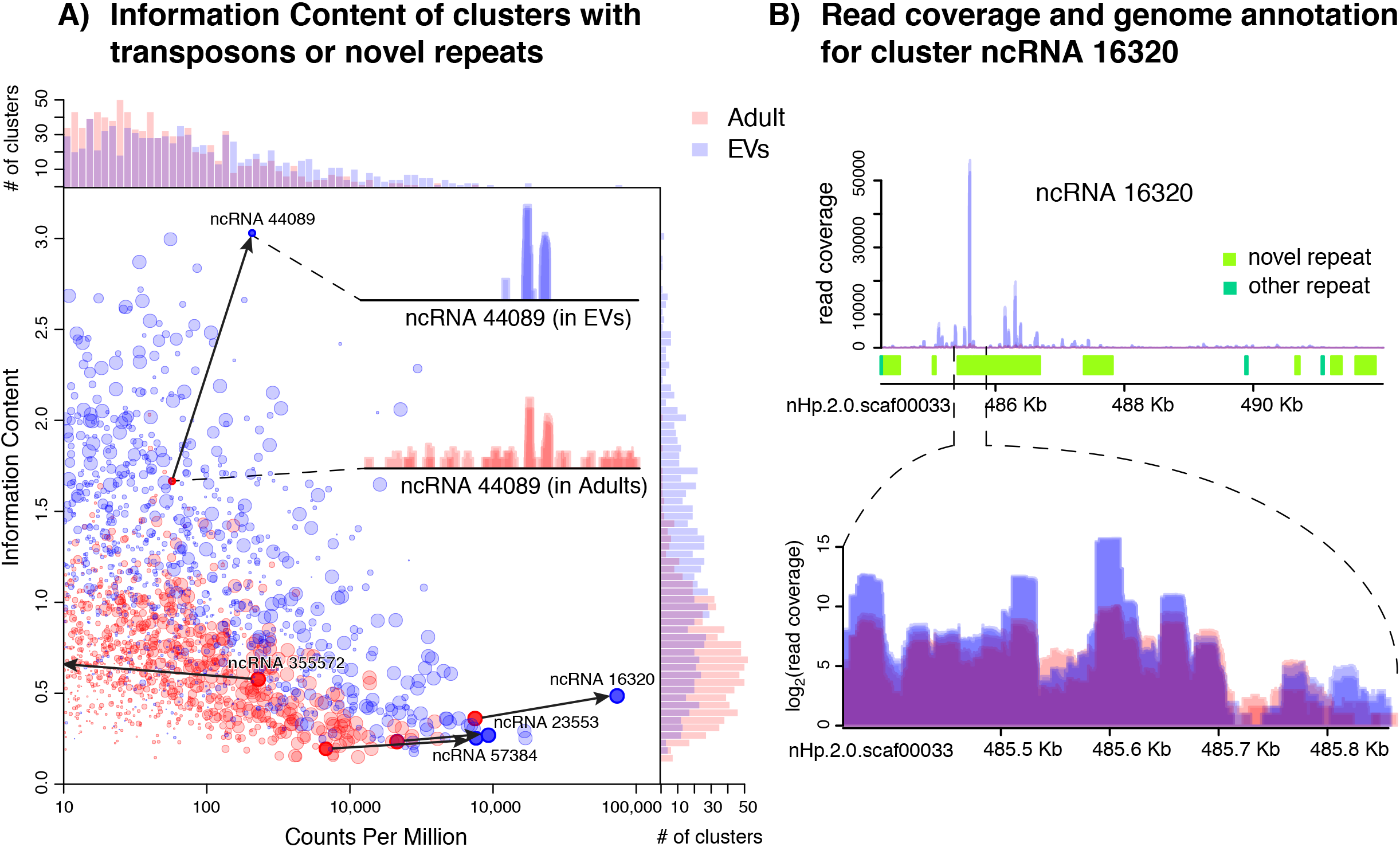
Clusters with transposons or novel repeats have higher Information Content in extracellular vesicles than in adults. Dot plot comparing Counts Per Million and Information Content of all clusters with transposons or novel repeats (A). Top and side barplots show the number of clusters at each value of the X or Y-axis respectively. Inset: example of read coverage for cluster ncRNA_44089. Read coverage (top), annotation of repeat elements (middle) and zoomed-in read coverage (bottom, in log_2_-scale to distinguish individual libraries) for the most highly expressed cluster in EVs (B). In all cases blue indicates EV and red indicates Adult libraries.

To determine whether exWAGO specifically associates with the EV-enriched sRNAs produced from novel repeats, we immunoprecipitated the adult nematode lysates using an antibody raised against exWAGO and analysed by qRT-PCR the co-purified sRNAs. The siRNAs that derive from EV-enriched clusters immunopurified with exWAGO and were depleted in the unbound fraction (Figure 6). We observed the opposite pattern with the IgG bead control. In contrast, siRNAs derived from selected clusters that are abundant in adults but not represented in the EVs did not copurify with exWAGO, nor did Y RNAs or a conserved miRNA (Figure 6). To confirm that selective export of these siRNAs by exWAGO, we carried out an immunoprecipitation of exWAGO from *H. bakeri* EVs. As shown in Figure 6C, the three siRNAs that are enriched in EVs co-purify with exWAGO (~9 fold enrichment for siRNAs from loci 16320 and 57384 and 1.5 fold enrichment for siRNA from loci 23553) whereas the Y RNA and miR-100 do not. These results suggest that specific siRNA sequences bind to exWAGO, which mediates their encapsulation in EVs and defines the population of vesicular sRNAs that are secreted into the host environment.

**Figure 6:**
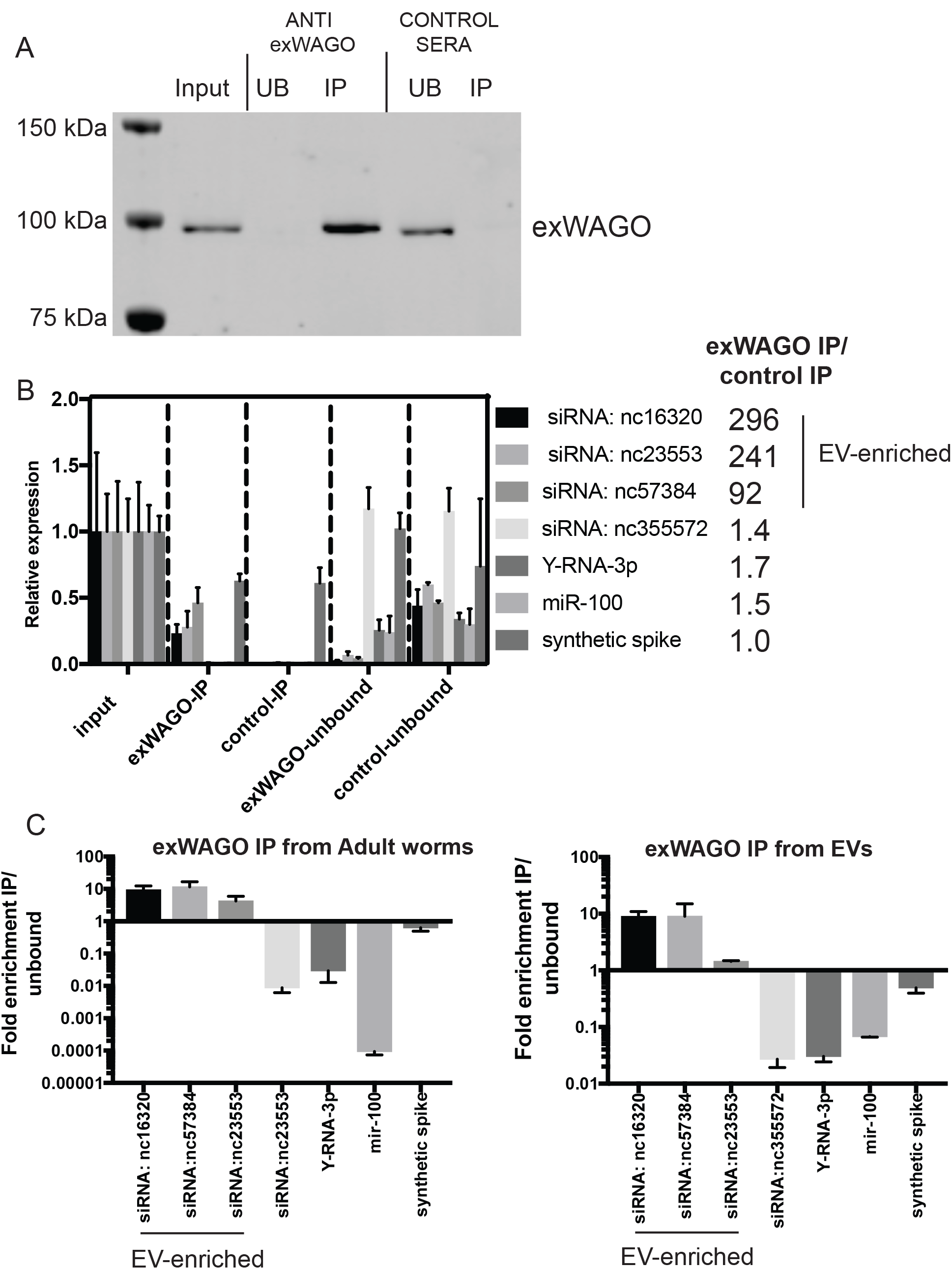
Immunoprecipitation of exWAGO and detection of associated RNAs. A) Western blot to detect exWAGO following immunoprecipitation of 10 ug adult worm lysates with exWAGO anti sera or control (naïve) sera. Equivalent volumes input and unbound were loaded (unbound is defined as first flow-through from beads). B) qRT-PCR analysis of samples from (A) for siRNAs derived from EV-enriched or adult-enriched clusters as well as Y-RNA and miR-100 (n=3, mean and standard deviation shown). Ratio of indicated RNAs in IP with exWAGO anti sera versus control (naïve) sera is noted on right. C) Fraction of indicated RNAs in exWAGO IP product versus unbound (flow through) from adult worm IPs (left), data as in (B) compared to IPs directly from 16 ug purified EVs (right) n=2. Mean and standard deviations are shown. To control for recovery of RNA during extraction, a synthetic spike was included prior to extraction which varied <2 fold across all sample types.

## Discussion

That small RNAs are transferred within organisms, and between organisms, has many implications in cross-species communication and disease. However, there are many questions regarding how RNAs are selected for export from the donor, which RNAs are transferred to the recipient, and whether and how these RNAs function within the recipient. Here we have examined the question of the specificity of packaging of sRNAs in EVs in the model parasite *H. bakeri* using comparative analyses of the origin of sRNAs within the body of the nematode *versus* those selected to be exported in EVs. We compare this to evolutionary analyses of the sRNA machinery in the parasite and the closely related, free-living *C. elegans*. We find that secondary siRNAs within adults of both the free-living and the parasitic nematodes are largely produced to target mRNAs and retrotransposons by antisense pairing along their entire length, and are therefore associated with endogenous gene regulation and defence. In contrast the siRNAs within EVs secreted by the parasite do not appear to be a stochastic sampling of those detected in the adult nematodes but are specifically enriched for those produced from transposons and newly-evolved, repetitive regions in the genome. We do not see this pattern of enrichment with siRNAs detected in the non-vesicular fractions of the secretion product. This suggests both molecular and evolutionary selectivity in what is packaged in EVs.

We identify exWAGO as one mechanism of selective siRNA encapsulation and export in the nematode parasites. Our immunoprecipitation experiments with exWAGO demonstrate association with EV-enriched siRNAs but not Y RNAs or miRNAs in EVs (6). Mechanistically, we envision three processes that could contribute to the total RNA present in EVs, acting independently or together. The EVs could be passively loaded with the sRNAs present in the cell type from which EVs are exported. It is likely that EVs are released from the intestine (6). Secondly, the sRNAs could be actively loaded by some intrinsic property, perhaps related to their specific biogenesis pathway. Lastly, the sRNAs could associate with a specific RNA-binding protein, as has been shown in some mammalian systems (8-10). Our immunoprecipitation data suggest that associative binding occurs for the siRNAs and we identify exWAGO as the mediator of this selective export. Intriguingly exWAGO is highly conserved and dominant in all Clade V parasitic nematodes examined, apart from *D. viviparus*, and we have further shown that it is also secreted in the rodent parasitic nematode *N. brasiliensis*. We propose therefore that the mechanism of exWAGO-mediated siRNA export extends beyond the *H. bakeri* model. At the same time, we cannot rule out that exWAGO plays additional functions inside the parasite.

The sRNAs selected for export with exWAGO derive from regions of the *H. bakeri* genome that are repetitive and novel, which may reflect recent, dynamic evolution of this putative host manipulation system. Rather than derive sRNAs from conserved loci, and risk self-directed effects, selection may exploit the rapidly evolving non-genic portion of the genome to generate evolutionarily novel but host-relevant sRNA loci. It will be informative to understand whether and how these associate with host genes inside host cells, since we know the EVs can be internalized by host cells (6,37).

We identified the SAGO and PPW proteins in *C. elegans* and related *Caenorhabditis* species as diverged exWAGO orthologues. Although as yet we have no evidence that the SAGOs (or any other AGOs) are exported extra-organismally from *C. elegans*, preliminary data suggest these may both have common localization in the intestine ((6) and Claycomb, Seroussi, unpublished). It will be of interest to understand whether SAGO and PPW function within *C. elegans* can shed light on the roles of exWAGO.

Very little is understood regarding the evolution of cross-species communication. That pathogens use small RNAs to modulate their hosts is not unexpected, as it has been well documented in interactions between parasitic fungi and plants (38) and it is similar, conceptually, to the evolution of miRNAs in certain viruses (39). In contrast to viruses, however, extracellular parasites such as *H. bakeri* require a mechanism for transporting specific sRNAs into host (mammalian) cells. The packaging of siRNAs in EVs by exWAGO provides a key mechanism for selectivity in export. EVs have also recently been implicated in the transfer of RNA in plants, in this case from the plant cells to fungal parasites although selectivity mechanisms are not known (40). We do not yet know if exWAGO is solely involved in export, or is also involved in mediating functional effects inside the recipient cells. We also do not yet know if the non-vesicular exWAGO plays any functional role. Several studies suggest mammalian Argonautes can be highly stable in extracellular environments, outside of EVs, but whether and how these are internalized by cells is not known.

Further work is required to understand the individual and collective contributions of all of the EV cargos in host cell modulation. This work establishes a parasite Argonaute as a sorting mechanism for selective export of siRNAs in EVs, indicates that focusing only on miRNAs can be misleading in some contexts and provides an important framework for interrogating new parasite-host interactions and their consequences on infection.

## AVAILABILITY

The new genome assembly and annotation for *Heligmosomoides bakeri* are available through WormBase ParaSite (http://parasite.wormbase.org/Heligmosomoides_polygyrus_prjeb15396) and the raw PacBio sequencing data are available at NCBI (https://www.ncbi.nlm.nih.gov/bioproiect/PRJEB15396).

## ACCESSION NUMBERS

The raw and processed small RNA sequencing files are available through GEO (accession number GSE117169).

## ACKNOWLEDGEMENTS

We thank Tuhin Maity for preparation of *C. elegans* samples and Rick Maizels for ES from *N. brasiliensis*.

## FUNDING

This work was supported by HFSP grant RGY0069 to AHB, CA-G and JMC.

## CONFLICT OF INTERESTS

The authors declare no competing interests.

## Notes

#### Summary of Updates

This is a revised version of the manuscript following peer review and including new data in Figure 6 and Supplementary Information.

