## Supplementary Material for "Secretion of an Argonaute protein by a parasitic nematode and the evolution of its siRNA guides"

**Supplementary Methods**

**Supplementary Table 1**: New *H. bakeri* genome assembly information.

**Supplementary Table 2:** Proteomes included in the OrthoFinder clustering analysis.

**Supplementary Figure 1: Quantification of exWAGO copy number in extracellular vesicles.** A) Western blot of ultracentrifuge pellet and supernatant of *H. bakeri* secretion product, using an antibody against exWAGO. Total protein amounts loaded are indicated. B) Relative quantification of western blot shown in (A) using mean band intensities with Image Studio Lite (LI-COR). C) Western blot of biological replicates of sucrose gradient purified EVs alongside two sets of recombinant Flag-His-tagged exWAGO protein. The *notes the mobility of the native exWAGO and ** is the mobility of the tagged version. Ladders are noted with "L1" and "L2" and the size of each band noted on each side of the blot. D) The band intensities of two sets of recombinant exWAGO and five replicates of known *H. bakeri* EV number are shown.

**Supplementary Figure 2: Conservation of RNAi pathway in Clade V**

Presence or absence of orthologues to *C. elegans* genes associated with RNAi pathways in Clade V organisms. Phylogenetic relationship is shown at the top of table, gene identities across each species are detailed in https://github.com/DRL/chow2018.

**Supplementary Figure 3:** Expression of all sRNA clusters (dots) comparing the average counts-per-million (X-axis) to the fold-change of polyphosphatase-treated relative to untreated libraries (Y-axis). Blue and red dots highlight those clusters identified respectively as polyP (enriched in polyphosphatase treated libraries) or monoP (similar normalised expression between treated and untreated libraries) for *C. elegans* adult nematodes (A), and *H. bakeri* adults (B) or extracellular vesicles (C). Clusters containing known miRNAs (expected to be monoP) are highlighted in gold. The numbers in parenthesis in the top-right labels of each figure represent the number of clusters defined for each category (polyP, monoP and miRNAs).

**Supplementary Figure 4:** Distribution of reads in annotated categories for each type of library, comparing the reads falling within monoP or polyP clusters to all reads. Plots are shown for untreated libraries for *C. elegans* (A) and *H. bakeri* (B), and polyphosphatase-treated libraries for *C. elegans* (C) and *H. bakeri* (D).

**Supplementary Figure 5:** Comparison of *C. elegans* and *H. bakeri* genome sizes and fractions of each genome devoted to annotated.

**Supplementary Figure 6: Distinct siRNA populations in *H. bakeri* EVs compared to the non-vesicular secretion product**. A) Multi-Dimensional Scaling plot showing the relationship between adult, supernatant and EV libraries, quantifying all the 20-25nt reads that fall within polyP-enriched clusters. B) Line plot showing the average percentage of these reads in each annotation category, for the adult, supernatant and EV libraries. C) qRT-PCR analysis of relative fraction of EV-enriched siRNAs, adult-enriched siRNA, Y RNA and miRNA levels in EVs compared to ultracentrifuge supernatant, mean and standard deviation shown (n=2).

#### Supplementary Methods

##### DNA extraction & Sequencing

To extract genomic DNA from adult *H. bakeri*, the nematodes were washed twice with sterile PBS and re-suspended in Puregene cell lysis buffer (Qiagen) before being triturated by hand using a sterile mortar and pestle under liquid nitrogen. The ground nematode extract was thawed and digested with 100 µg proteinase K (Qiagen) at 65°C with gentle shaking overnight. RNA was removed by subsequent digestion at 37°C for 1hr with RNase A (100 µg; Qiagen). Puregene Protein Precipitation Solution (Qiagen) was added and the digests incubated on ice for 5 min. Precipitate was removed by centrifugation at 13,000 rpm at 4°C for 10 min, and the supernatant transferred to a new tube. Isopropanol was added to the supernatant and samples placed at −20°C for at least 1 hr, followed by centrifugation and ethanol precipitation. Re-suspended DNA was treated with 5 µl RiboShredder mix (Epicentre) at 37°C for 2 hr prior to subsequent purification with the Zymo Research Genomic DNA Clean & Concentrator kit following manufacturer's instructions. Genomic DNA integrity and molecular weight were verified by agarose gel electrophoresis and two template libraries with 10 kb inserts were prepared. PacBio sequencing was carried out on the PacBio RS II platform following the standard protocol with a C2 sequencing kit at the 1×120-min acquisition mode (Centre for Genomic Research at University of Liverpool, UK). The run was carried out with diffusion-based loading and analysed using standard PacBio primary data analysis. Illumina short-read data were generated from independent DNA preparations, using 350 base and 550 base insert libraries on a HiSeq2500 instrument (paired end 125 base reads) (Edinburgh Genomics at University of Edinburgh, UK).

##### RNA preparation from adult worms and RNA sequencing

*C. elegans* cultures were grown at a density of 100,000 worms per plate, on 150mm plates with 2ml of 5X concentrated OP50 to the gravid adult stage (containing embryos; 65h post L1 at 20°C), and harvested and flash frozen as in (1). *H. bakeri* were collected as above and disrupted in 700 µl of Qiazol (Qiagen) using mechanical disruption with 5 mm stainless steel beads (Qiagen) on a Tissue Lyser II (30hz for 2 min twice; Qiagen). Total RNA was extracted using a miRNAeasy mini kit (Qiagen) following the manufacturer's instruction. RNA was treated with Turbo DNA-free kit (Thermo Fisher) to remove residual DNA. Total

RNA was treated with RNA 5' Polyphosphatase (Epicenter) following manufacturer's instructions, before library preparation. Libraries for small RNA sequencing were prepared using the CleanTag small RNA library prep kit (Trilink) according to the manufacturer's instructions, using total RNA from adult worms (30 ng), and sucrose-gradient purified EVs (equivalent $1X10^{10}$ EVs measured by Nanosight, Malvern). For all samples, 1:12 dilutions of both adapters were used with 18 amplification cycles (TriLink BioTechnologies). The products between 140-170bp were size selected and sequenced on an Illumina HiSeq high output v4 50bp SE in Edinburgh Genomics. For stranded mRNA sequencing, the Illumina TruSeq was carried out by Edinburgh Genomics and the libraries sequenced on an MiSeq (reagent Kit v2) with 100PE in Edinburgh Genomics.

**Genome assembly and annotation**

The quality of Illumina reads was checked with FastQC (http://www.bioinformatics.babraham.ac.uk/projects/fastqc). Adapters were removed with Cutadapt (2), low-quality bases were trimmed with Trimmomatic (3), and reads were error-corrected with BLESS (4) using a kmer size of 21. A preliminary assembly was performed with Velvet (5) and library insert sizes confirmed after read mapping using bowtie2 (6). The Velvet assembly was inspected for contamination with blobtools (7), with no contamination being detected. The short Illumina data were assembled and gapfilled using Platanus (8), and scaffolded using transcriptome evidence (described below) with SCUBAT2 (Koutsovoulos G. SCUBAT2. https://github.com/GDKO/SCUBAT2) Long-read PacBio data (a total of 10.2 Gb from reads with an N50 of 9,411 bases) were used to further scaffold and gapfill the assembly with PBJelly (9). RNA-Seq reads were assembled with Trinity (10) resulting in 38,777 transcripts in 30,483 gene components. The transcriptome assembly was filtered based on expression (>1 TPM) and isoform percentage (>0%) calculated by kallisto (11), reducing the dataset to 32,595 transcripts. Transcripts that could encode a protein with at least 50 amino acids were selected to use with SCUBAT2. RepeatModeler (Smit, AFA, Hubley, R. RepeatModeler http://www.repeatmasker.org) and RepeatMasker (Smit, AFA, Hubley, R & Green, P. RepeatMasker http://www.repeatmasker.org) were used to identify and mask repeated regions in the genome prior to gene finding. We used the BRAKER (12) pipeline to predict protein-coding genes using the RNA-Seq reads as evidence. We combined the BRAKER general feature format (gff) file and the transcriptome assembly within MAKER2 (13) to predict untranslated regions of transcripts (UTRs) and remove low quality gene predictions. This curated set was used for downstream analyses.

**Identification of *H. bakeri* exWAGO orthologues in other Rhabditomorpha**

Loci orthologous to the *H. bakeri* secreted WAGO (exWAGO) were identified using BLAST (14) searches of the genome-derived proteomes of *Haemonchus contortus*, *Necator americanus* and *Pristionchus pacificus*. The gene model of each identified orthologue was then evaluated with RNA-Seq data from the relevant species, and corrected if necessary. Using an alignment of these four proteins, a custom hidden Markov model (HMM) was constructed to identify homologues in other nematode genomes using HMMer (15) within GenePS (Koutsovoulos G https://github.com/jgraveme/GenePS). The discovered gene models were corrected based on alignment of the HMM profile, *de novo* Augustus (16) prediction, the previous predicted gene models and RNA-Seq data if available. Orthologues in *Caenorhabditis* species were validated through analysis of reciprocal best BLAST matches. Protein sequences of exWAGO orthologues were aligned with MAFFT (17) and the alignment was analysed with PHYML (using the LG+G model) (18,19). Bootstrap support was calculated from 100 bootstrap replicates.

**Ortholog clustering**

Protein sequences of 21 nematode species were retrieved from the sources specified in Supplementary Table 2. Sequences below a length of 30 residues and containing more than one non-terminal stop codon were removed. Manually annotated *H. bakeri* exWAGO orthologues were added to the corresponding proteomes. Sequence similarity searches were performed using BLAST v2.4.0+ (-evalue 1e-5 -outfmt '6' -seg yes -soft_masking true -use_sw_tback). Protein clustering was carried out using OrthoFinder v1.1.4 (20) under the MCL inflation value of 3.0. Functional annotation of proteins was carried out via InterProScan v5.22-61.0 (21) against PFAM v30.0 (22) and SignalP-Euk v4.1 (23). Orthogroups were analysed using KinFin v1.0.3 (24) by providing functional annotation and the phylogenetic tree of the taxa. The orthogroups were screened based on previously described *C. elegans* WAGOs (25) using the KinFin script get_count_matrix.py. Output files are deposited at https://github.com/DRL/chow2018.

**Analysis of Argonaute expression levels in Clade V**

To quantify the overall Argonaute transcript levels in Clade V parasitic nematodes, we assessed the NCBI/SRA (https://www.ncbi.nlm.nih.gov/sra) and downloaded the available Illumina raw data of mixed adults or female adults life-stages. There are 13 paired-end libraries corresponding to six nematodes (including *H. bakeri*), detailed in Supplementary Table 3. The fastq-dump from the SRA toolkit (NCBI 2011) was used to obtain sequence files in FASTQ format. Sequences were filtered for low quality reads and adapters using Trim Galore v0.4.5 (https://www.bioinformatics.babraham.ac.uk/projects/trim_galore/). Transcripts were then quantified using corresponding reference transcriptome from WormBase Parasite (26) within Salmon v0.10.1 (27). The mean tpm (n=2) of *C. elegans* Argonaute transcripts were obtained from EMBL-EBI Expression Atlas (28).

**Proteomics of excretory-secretory products**

Total protein from three replicates *N. brasiliensis* ES (2.5-5ug EVs and 5 ug supernatant) was loaded on a 4-12% Tris-Bis NuPAGE gel (Invitrogen) and electrophoresed before overnight in-gel digestion as described (29). Peptide extracts were dried by Speedvac and the dried peptide samples were re-suspended in MS-loading buffer (0.5% trifluoroacetic acid in water) then filtered before HPLC-MS analysis. Analysis was performed using an online system of a nano-HPLC (Dionex Ultimate 3000 RSLC, Thermo-Fisher) coupled to a QExactive mass spectrometer (Thermo-Fisher) with a 300µm x 5mm pre-column (Acclaim Pepmap, 5µm particle size) joined with a 75µm x 50cm column (Acclaim Pepmap, 3µm particle size). Peptides were separated using a multi-step gradient of 2–98% buffer B (80% acetonitrile and 0.1% formic acid) at a flow rate of 300 nL/min over 90 min. Data from MS/MS spectra was searched using MASCOT against a *N. brasiliensis* databases (WormBase Parasite). The parameters used in each search were: (i) missed cut = 2, (ii) fixed cysteine carbamidomethylation modification, (iii) variable methionine oxidation modification, (iv) peptide mass tolerance of 10ppm, (v) fragment mass tolerance of 0.05 Da. Search results were exported using a significance threshold (p-value) of less than 0.05 and a peptide score cut off of 20.

**Nanoparticle tracking analysis (NTA)**

NTA was carried out using a NanoSight LM14 instrument (Malvern Instruments, Malvern, UK). Vesicles were diluted with 0.1 µm-filtered PBS prior to analysis. NTA 2.2 software was used to record and analyse the samples. The camera level was set to 15 and the detection threshold to 5. Minimum expected particle size, blur and minimum track length were set to auto. The background extraction was set to On.

Three measurements with 60s recording were taken for each sample. The mean and standard error of the mean (SEM) were calculated and plotted using GraphPad Prism 7 (GraphPad Software).

**Transmission electron microscopy (TEM)**

For visualization of the vesicles, the purified *H. bakeri* EVs were fixed in 2% paraformaldehyde (PFA), deposited on Formvar-carbon-coated EM grids and treated with glutaraldehyde before treatment with uranyl oxalate and methyl cellulose as described elsewhere previously (30) and then viewed in a Philips CM120 TEM. Images were taken on a Gatan Orius CCD camera.

**Annotation of known families of ncRNA in *H. bakeri* by homology**

To expand the annotation of *H. bakeri* into the non-coding realm, we predicted ncRNA families with Infernal 1.1.1 (31) using covariance models from Rfam 12.0 (32). For transfer RNA annotation, we used tRNAscan-SE 1.3.1 (33), and RNAmmer 1.2 (34) for ribosomal RNA. We downloaded all mature sequences from miRBase 21 (35) to annotate known miRNAs with MapMi 1.5.9 (36). The sequences of predicted and curated miRNAs and piRNAs from a previous publication (30) were also transferred to the new genome using BLAST and requiring perfect hits. yRNA annotation is based on our previous report (30).

**Annotation of ncRNA in *H. bakeri* using sRNA-seq data**

We also used all our sRNA-seq results to predict novel ncRNA producing regions. For this, we used two different approaches: ShortStack 3.8.3 (37) and miRDeep2 (38). For mapping and analysis by ShortStack, we combined all libraries for each genome as a single input, and used the following parameters: --pad 10, --mincov 10, --dicermin 18, --dicermax 32. ShortStack will thus predict regions or clusters on the genome where many sRNAs are mapped. We also used miRDeep2 to discover new miRNAs. For the miRDeep2 alignment procedure we used the combination of all libraries and parameters: -c -j -l 18 –m -q. For the miRDeep identification procedure, we used parameters: -a 100, -g 500. As input sequences, we used the high confidence *H. bakeri* mature sequences from our previous publication (30) and all mature and hairpin sequences from miRBase 21 (35).

**Annotation segmentation of the genome**

In order to understand how different parts of the genome led to sRNA production, we produced a non-overlapping segmentation of the genomic annotation. At the end, every base of the genome was assigned to a single type of annotation. For bases that had more than one type of overlapping annotation, we defined a simple hierarchy to choose the preferred one. The hierarchy consisted of: miRNA > yRNA > piRNA > tRNA > rRNA > snRNA > snoRNA > other_ncRNA > known retrotransposons > known transposons > mRNA exons > mRNA introns > satellite repeats > novel repeats > low complexity and simple repeats. For example, if a base overlapped with a miRNA and an intron on the same strand, it was assigned as a miRNA. Different annotation was allowed on either strand, so a single base pair could be assigned as an intron on one strand, and as a miRNA on the other strand. This was accomplished within R, extensively using objects and functions from the GenomicRanges package (39). The total number of bases assigned to each annotation type in both genomes is presented in Table 1. For all analyses that required defining a single annotation for each read (Figure 4), we used the annotation of the base aligned to the central position of the read.

**Definition and classification of sRNA-producing clusters in the genome**

To define discrete sRNA-producing regions in the genome, we selected ShortStack clusters as our main reference loci. Some of these clusters overlapped exon-intron boundaries. Since we were interested in distinguishing the sRNA reads coming from introns (possible degradation products) and exons (potential siRNAs), we split all overlapping clusters at exon-intron boundaries. A total of 417,292 clusters were defined for the *H. bakeri* genome, and 103,278 for *C. elegans*.

**Expression quantification of sRNA-producing clusters**

In order to obtain a level of expression for each sRNA-producing cluster, we counted mapped reads using the *findOverlaps* function from GenomicRanges R package (39) with parameters minoverlap=16 and ignore.strand=TRUE. With this we obtained count tables, where each cluster was a row and each library a column. We also calculated coverage mapping to clusters using the *coverage* function from GenomicRanges R package using counts as weights.

**Differential expression analysis of sRNA-producing clusters**

Exploratory analysis of the expression levels of all clusters across libraries, led us to the conclusion that we had two distinct types of clusters: those producing mostly sRNAs with a 5' mono-phosphate (expressed in both types of libraries) and those producing mostly sRNAs with a 5' poly-phosphate (expressed mostly in the polyphosphatase-treated libraries). In order to consistently predict these two types of clusters, we performed differential expression analyses using the edgeR package (40). *C. elegans* libraries were analysed on their own, and since *H. bakeri* adult and EV libraries were quite distinct (according to MDS plots), they were analysed separately. We first imported the count tables, chose the desired libraries, and selected only clusters with at least 0.5 counts-per-million in at least 2 libraries. Next, we performed an edgeR analysis with TMM normalisation and using only the estimated common dispersion. To define the monoP-enriched clusters (with higher relative expression in untreated libraries, since untreated libraries do not contain 5' poly-phosphate reads) we performed a *glmTreat* test, searching for clusters that were significantly more abundant in the untreated libraries. We visually confirmed that the selected clusters contained the expected types of annotation (rRNA, tRNA, miRNA), and formed a consistently horizontal cloud in abundance vs fold-change (MA) plots. To define these monoP-enriched clusters, we used different cutoffs: for *C. elegans* a fold-change of 5 and False Discovery Rate (FDR) of 0.01, for *H. bakeri* adult libraries a fold-change of 2 and FDR of 0.01, for *H. bakeri* EV libraries a fold-change of 1.05 and FDR of 0.25. In all cases we calculated the FDR using the Benjamini-Hochberg method (41). Supplemental Figure 1 shows these results, highlighting the overlap with miRNAs. We next used the miRNA-containing monoP-enriched clusters as a baseline (since they should be equally expressed in untreated and treated libraries) to re-normalise the data. To define the polyP-enriched clusters, we again used a *glmTreat* test, searching for those with significantly more expression in the polyphosphatase-treated libraries. For the *C. elegans* and *H. bakeri* adult libraries we used a fold-change cutoff of 2 and 0.01 FDR, and for the EV libraries a fold-change of 1.5 and FDR of 0.2. The resulting polyP-enriched clusters are also indicated in Supplemental Figure 2.

**Information Content of sRNA-producing clusters**

To describe the pattern of reads covering each cluster, we calculated the coverage cluster Information Content (IC). IC is based on the coverage entropy of each cluster, compared to a uniform coverage distribution (maximum entropy) and is defined as *IC = log2(length(y)-entropy(y))*, where *y* is the cluster coverage. Entropy values were calculated with the *entropy* function from *entropy* R package (42). The total number of bases for each coverage value (discretized by the function y = *round(log10(coverage+1),1)*10*) were tabulated and converted into an entropy value. This was subtracted from the maximum entropy given the length of each cluster. For a perfectly uniform distribution (either no reads, or the same depth across the whole cluster, IC = 0. If IC > 0, there is more coverage variability, with the cluster presenting one or more “peaks”. To avoid biases caused by higher sequencing depth of the Adult libraries, we used the same number of amplification cycles for both types of libraries, and during the bioinformatic analysis we randomly sampled exactly 2.5 million mapped reads from two Adult and two EV libraries, before calculating the IC values as mentioned above.

**Argonaute Immunoprecipitations**

*H. bakeri* adult worms were lysed with worm lysis buffer (10 mM Tris-HCl, 150 mM NaCl, 0.5% NP40, 0.5 mM EDTA, cOmplete™ Protease Inhibitor Cocktail Tablets from Roche, pH 7) using mechanical disruption with 5 mm stainless steel beads (Qiagen) on a Tissue Lyser II (30hz for 2 min twice; Qiagen). The lysates were cleared by centrifugation (16,000×g) for 10 min at 4°C and immediately used for immunoprecipitation.. The supernatants (10ug each) were immunoprecipitated with rat polyclonal anti-exWAGO antibody (raised against full length protein) or rat normal IgG followed by protein L magnetic beads (Fisher Scientific). After immunoprecipitation, equivalent amounts (from 200 uL) of the input, flow-through and the immunoprecipitated product were kept for RNA extraction using the miRNeasy Serum/Plasma Kit (Qiagen). Briefly, a synthetic spike RNA (termed RT4) was added at 0.1pM to the Qiazol (5X sample volume) before RNA extraction as an internal control. The RNAs were eluted with 14ul RNase-free water. The small RNAs that were associated with *H. bakeri* exWAGO were analysed by qRT-PCR as below. The rabbit polyclonal anti-exWAGO antibody (as described above) was used for the western blot analysis of exWAGO.

**Detection of *H. bakeri* siRNAs by qRT-PCR**

For reverse transcription of RNA from exWAGO IPs, a fixed volume of 5 uL of total RNA was used as input (20 uL reaction volume) using the miScript RT II System (Qiagen) according to the manufacturer's protocol. cDNAs were diluted 10-fold in nuclease-free water for quantitative PCR. Quantitative PCR was carried out with the QuantiTec SYBR Green PCR kit (Qiagen), which includes a universal primer, according to the manufacturer's protocol. Primers for *H. bakeri* specific siRNAs and synthetic spike-in RNAs were used at a final concentration of 200 nM and were purchased from IDT and efficiency measured as in (43). Two technical replicates for each of the three biological replicates were included, as well as a nuclease-free water ("no template") control. The immunoprecipitated samples using exWAGO were defined as experimental group and those using rat normal IgG were defined as control group. The q-RT-PCR condition used was as follows: 1) pre-denaturation for 15 min at 95$^o$C, 2) 40 cycles of denaturation 15s at 94$^o$C, annealing 30s at 55$^{o}$C, and elongation 30s at 70$^{o}$C. Melting curve was included for each sample to check specificity. Fluorescence data collection was performed at the end of each annealing step. Data was collected on a Light Cycler 480 System (Roche) and Cq were calculated with the Light Cycler 480 software (version sw 1.5.1) with the "High Confidence" and "SYBR Green I / HRM Dye (465-510)" settings. No signals were detected in all NTC. Data were analysed using the $2^{-\Delta\Delta Ct}$ method described in (44).

List of qRT-PCR DNA primers used:

| | |
|---|---|
| EV-enriched_nc16320 | GATGACCAACCGGCTGTGGAAGC |
| EV-enriched_nc57384 | GTAGTTGGGGTGGTTGTAGG |
| EV-enriched_nc23553 | GAACGACTGCTTCTATGCCACCCGA |
| Adult-enriched_nc355572 | GGAACTCCCAACGGGCCCGGG |
| Y-RNA-3p | CGACAAAAGCTCGACCGGCGC |
| miR-100 | AACCCGTAGATCCGAACTTGTGT |
| Synthetic spike | CTTGCGCAGATAGTCGACACGA |

**Supplemental Table 1:** *Heligmosomoides bakeri* genome assembly

| **Feature** | ***Heligmosomoides bakeri*** **genome assembly v2.0** | ***Heligmosomoides bakeri*** **genome assembly v1.0** |
|---|---|---|
| Reference | This work | WTSI |
| Span (Mb) | 696 | 560 |
| G+C content (%) | 45.6 | 45.0 |
| Scaffold / contig N50 (kb) | 179.6 / 42.6 | 35.8 / 12.8 |
| Number of contigs | 23647 | 44728 |
| Genome CEGMA complete / partial (%) | 88.7 / 8.1 | 78.8 / 18.1 |
| Genome BUSCO (Nematoda) complete / partial (%) | 87.1 / 7.2 | 67.8 / 10.7 |
| Genome BUSCO (Eukaryota) complete / partial (%) | 87.8 / 1.7 | 74.3 / 8.9 |
| Transcriptome mapping | 96.3% | 72.3% |
| Number of protein-coding genes | 24371 | 27459 |

**Supplementary Table 2**: Proteomes included in the OrthoFinder clustering analysis. Prefix: prefix used for protein sequences. Species: species name. Source: source of protein files. WBPS8: WormBase ParaSite 8. EDI. CGP2:

| Prefix | Species | Source |
|---|---|---|
| ACEYL | *Ancylostoma ceylanicum* | WBPS8 PRJNA231479 |
| ACANT | *Angiostrongylus cantonensis* | WBPS8 PRJEB493 |
| ACOST | *Angiostrongylus costaricensis* | WBPS8 PRJEB494 |
| CAFRA | *Caenorhabditis afra* | CGP2 JU1286 |
| CBRIG | *Caenorhabditis briggsae* | WBPS8 PRJNA10731 |
| CCAST | *Caenorhabditis castelli* | CGP2 JU1956 |
| CELEG | *Caenorhabditis elegans* | WBPS8 PRJNA13758 |
| CSP1 | *Caenorhabditis monodelphis* | CGP2 JU1667 |
| CSP38 | *Caenorhabditis* sp. 38 | CGP2 JU2809 |
| CTROP | *Caenorhabditis tropicalis* | WBPS8 PRJNA53597 |
| DVIVI | *Dictyocaulus viviparus* | WBPS8 PRJEB5116 |
| HCONT | *Haemonchus contortus* | WBPS8 PRJEB506 |
| HPLAC | *haemonchus placei* | WBPS8 PRJEB509 |
| HPOLY | *Heligmosomoides polygyrus* | EDI v2 |
| HBACT | *Heterorhabditis bacteriophora* | WBPS8 PRJNA13977 |
| NAMER | *Necator americanus* | WBPS8 PRJNA72135 |
| NBRAS | *Nippostrongylus brasiliensis* | WBPS8 PRJEB511 |
| ODENT | *Oesophagostomum dentatum* | WBPS8 PRJNA72579 |
| OTIPU | *Oscheius tipulae* | EDI v2 |
| PPACI | *Pristionchus pacificus* | WBPS8 PRJNA12644 |

Caenorhabditis Genome Project *v2.*

#### Supplemental Figure 1

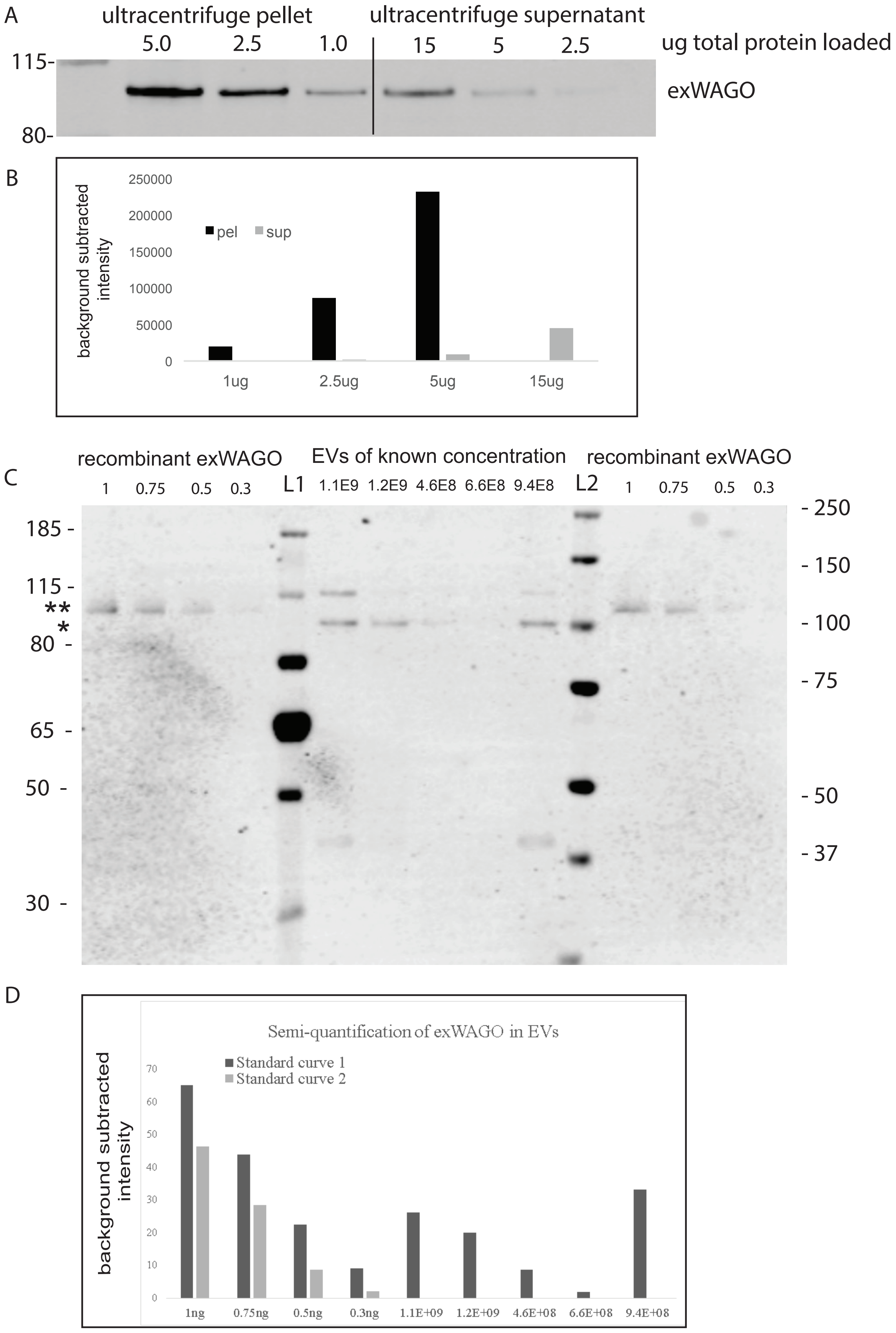

### Supplemental Figure 2

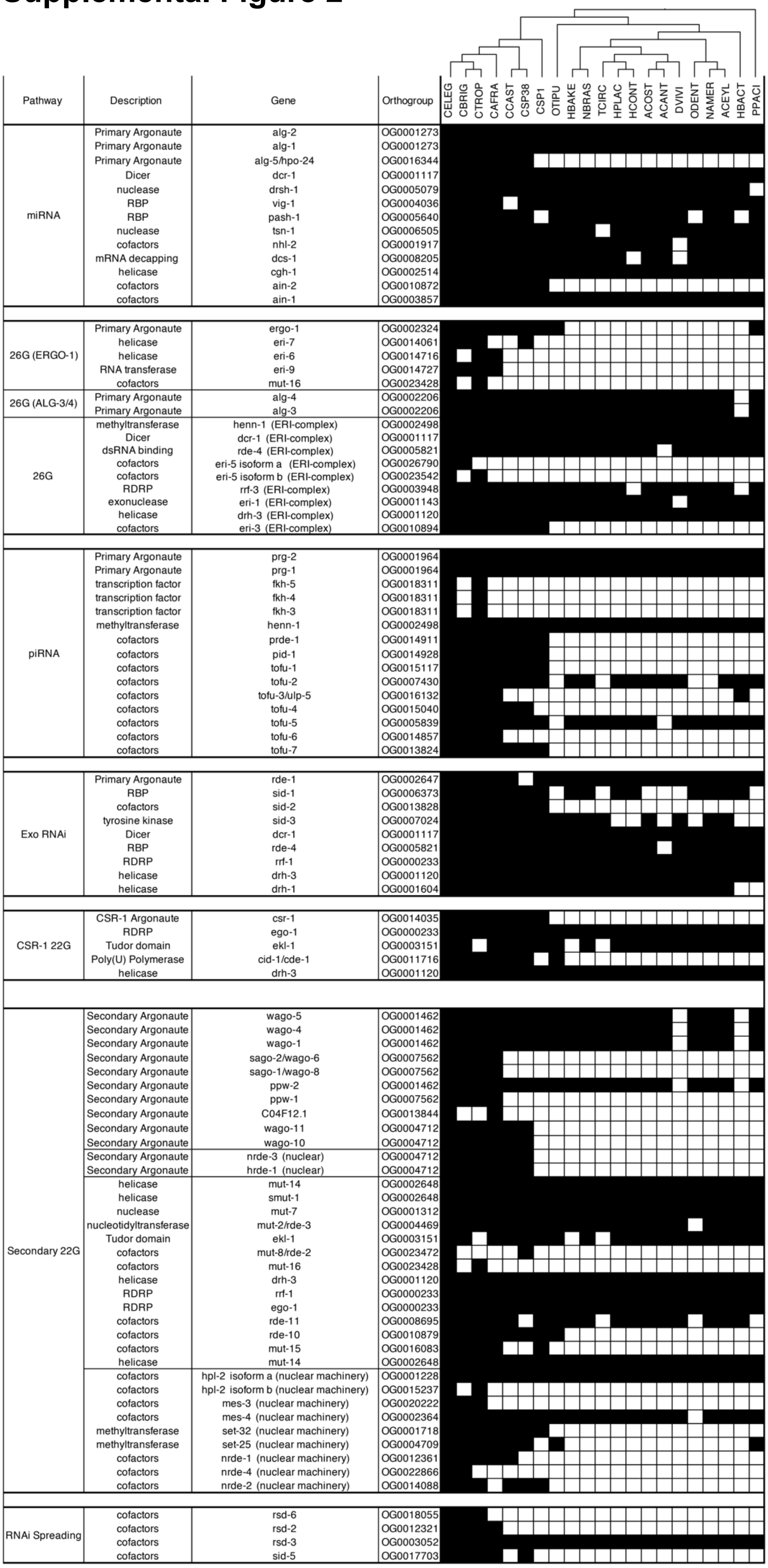

### Supplemental Figure 3

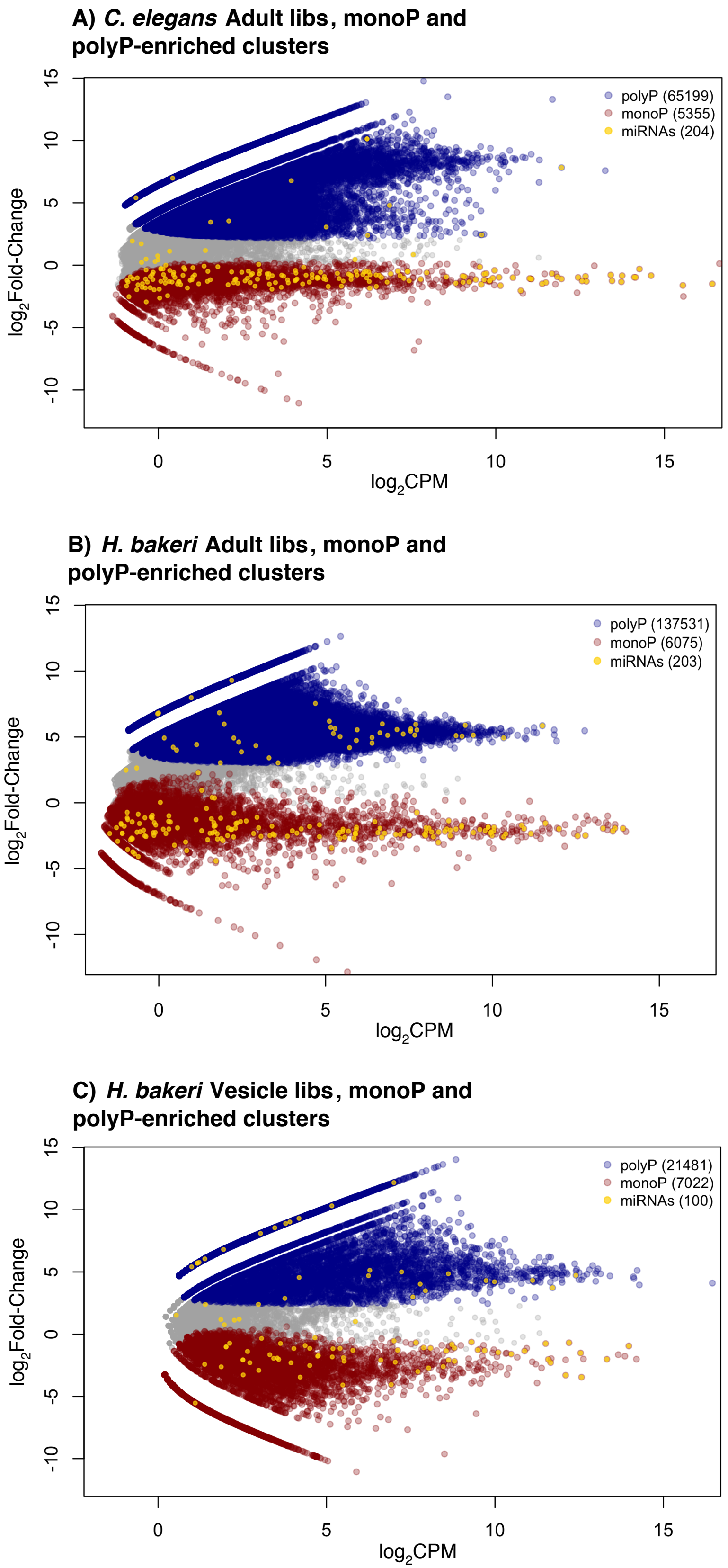

### Supplemental Figure 4

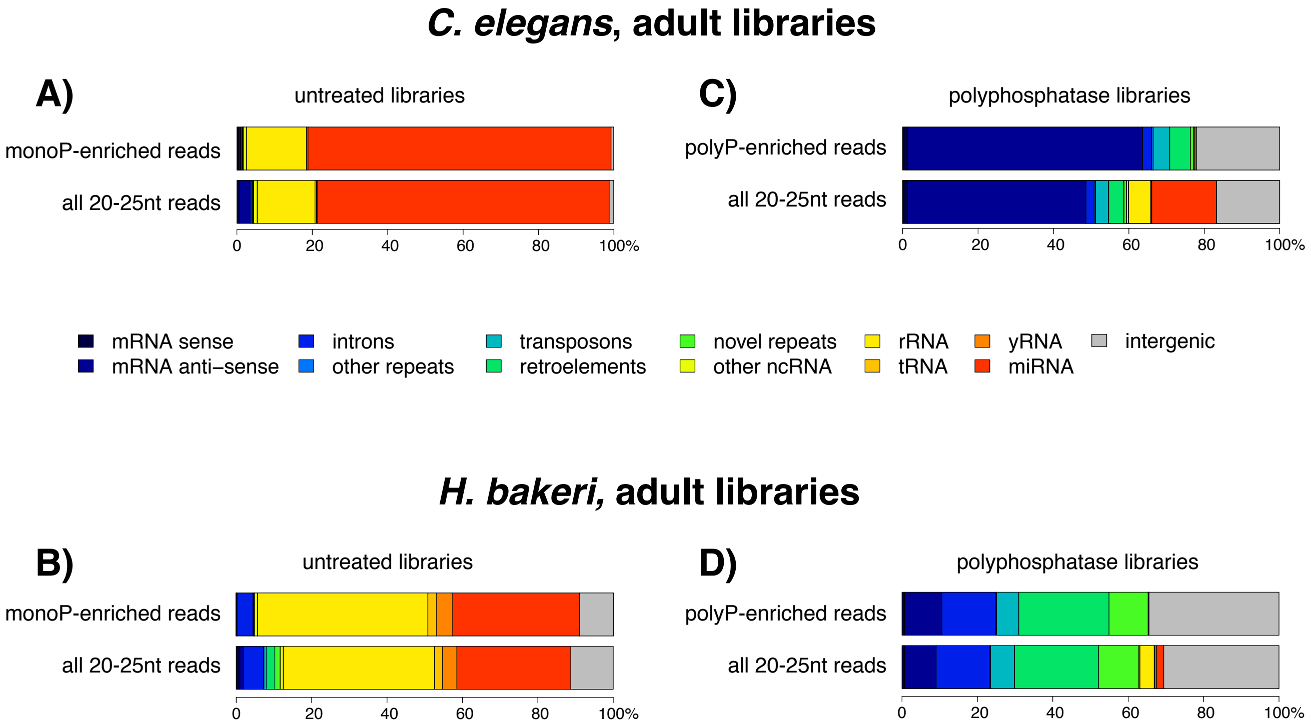

#### Supplemental Figure 5

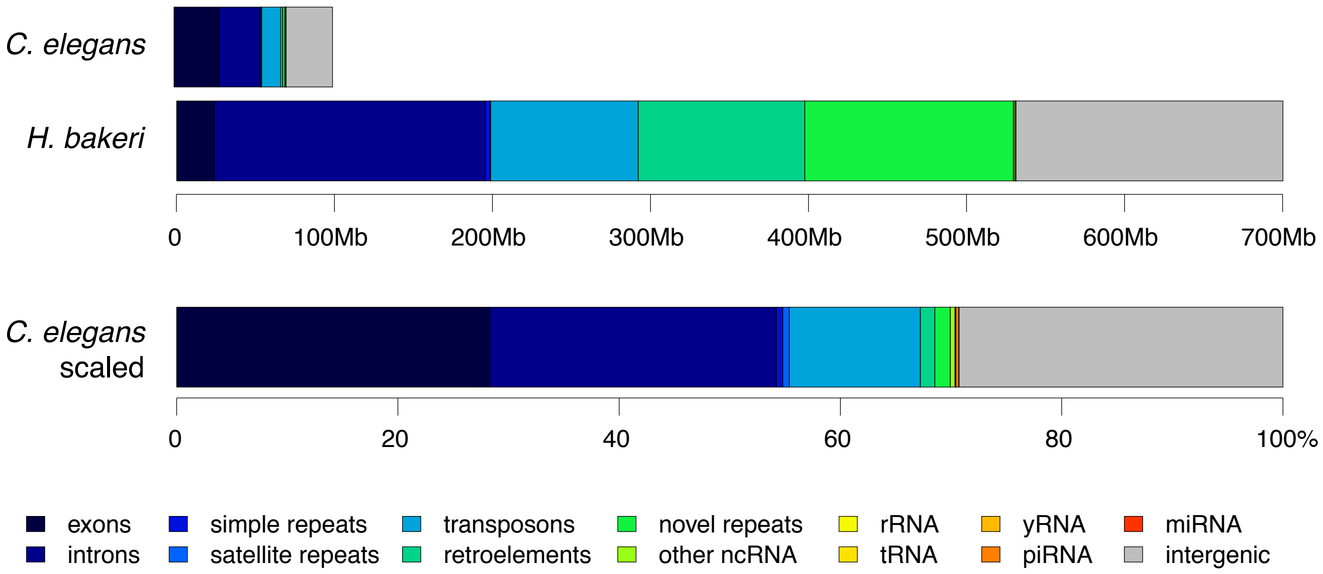

### Supplemental Figure 6

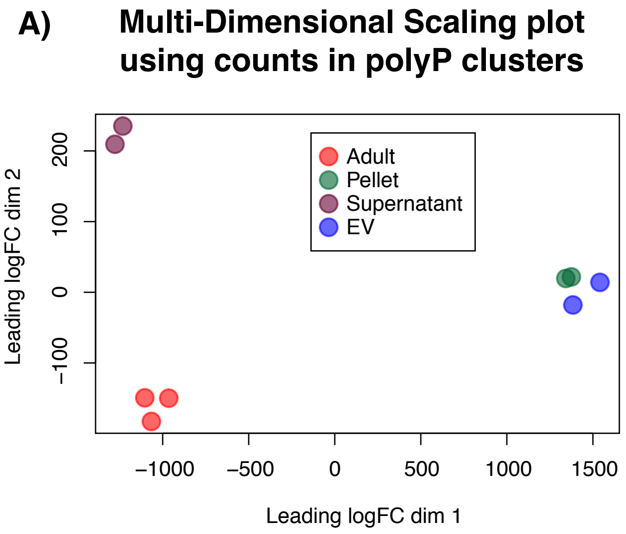


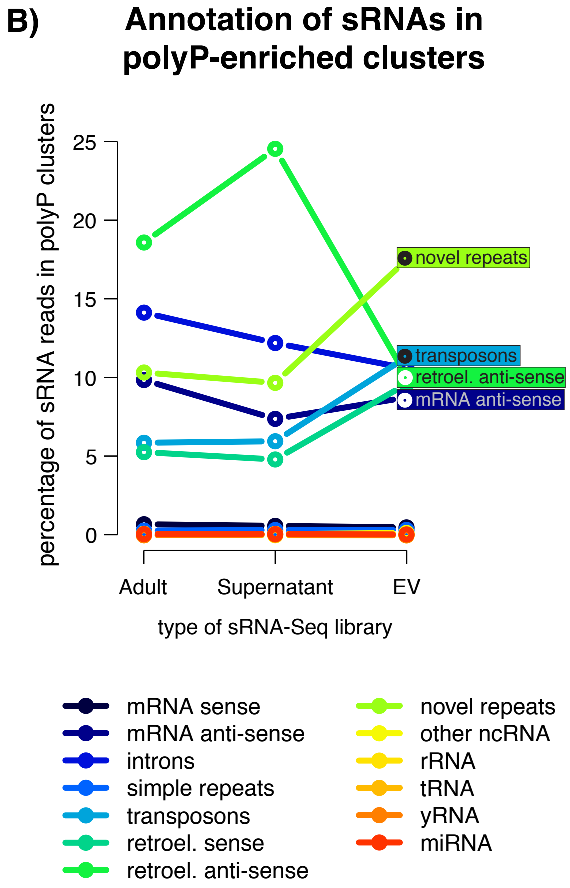


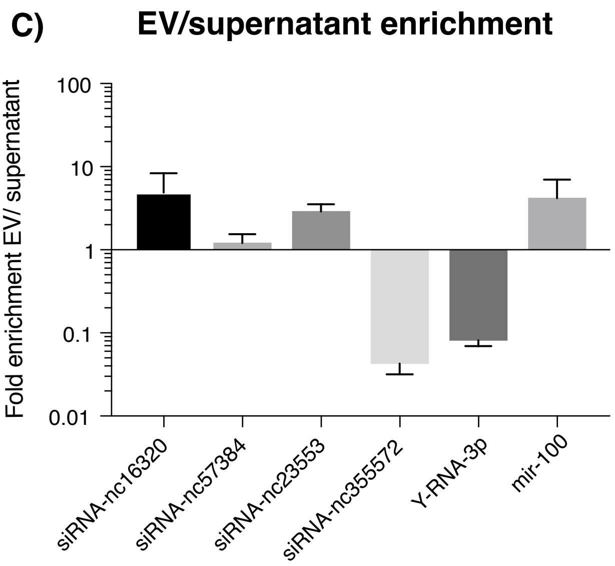
